# Local recurrence accounts for extended processing during occluded-object recognition

**DOI:** 10.64898/2026.09.10.750662

**Authors:** Mohammad Ahadzadeh, Karim Rajaei, Hamid Soltanian-Zadeh

## Abstract

Recognizing objects from incomplete visual input often requires processing beyond the initial feedforward sweep, but the relative contributions of local recurrence and long-range top-down feedback remain unclear. We combined source-localized magnetoencephalography (MEG), time-resolved decoding, backward masking, representational Granger causality, and computational modeling to examine these mechanisms during occluded-object recognition. We characterized neural dynamics in early visual cortex (V1-3), the lateral occipital complex (LOC), and inferotemporal-parahippocampal cortex (IT-PHC). Occlusion delayed the emergence of category information and elicited a late component that was selectively disrupted by backward masking, consistent with dependence on continued processing. These temporal changes occurred without detectable changes in the relative timing of regional responses or directed interareal interactions, including no measurable occlusion-related increase in feedback among the regions examined. To assess candidate computational mechanisms, we compared three nested model variants sharing a trained backbone: feedforward, local recurrent, and local recurrent with added long-range top-down feedback. Local recurrence improved recognition under occlusion and increased model-brain correspondence during the late, mask-sensitive interval, particularly in V1-3. Adding the implemented top-down pathway provided no consistent further benefit. Together, these findings favor prolonged local recurrent processing as an account of the additional computation supporting occluded-object recognition under the conditions tested.

## Introduction

Objects in natural scenes are often partially hidden by other objects. Recognizing them requires the visual system to extract identity from incomplete contours, surfaces, and contextual cues (Johnson & Olshausen, 2005; Kovacs et al., 1995; Tang et al., 2014). Occlusion poses a particular challenge because visible fragments must be assigned to the object or the occluder and integrated into a coherent percept (Kosai et al., 2014; Rensink & Enns, 1998). Studies of the neural and computational basis of this ability suggest that recognition can depend on processing beyond the initial feedforward response (Rajaei et al., 2019; Tang et al., 2018). Occluded-object recognition therefore provides a useful test case for determining how recurrent computation supports visual perception (Ernst et al., 2021; Wyatte et al., 2012).

Object recognition is commonly described as a hierarchical process in which information passes from early visual cortex to progressively higher-level regions of the ventral visual pathway (DiCarlo et al., 2012; DiCarlo & Cox, 2007). Feedforward models capture important aspects of the rapid emergence of object representations under relatively clear viewing conditions (Cadieu et al., 2014; Khaligh-Razavi & Kriegeskorte, 2014; Kubilius et al., 2019; Yamins et al., 2014; Yamins & DiCarlo, 2016). However, a single feedforward sweep may be insufficient when sensory evidence is ambiguous or incomplete (Ghodrati et al., 2014; Maniquet et al., 2025; Rajaei et al., 2019; Wyatte et al., 2014). Under these conditions, recurrent interactions may refine or sustain representations through repeated processing within cortical areas and reciprocal communication between hierarchical levels (V. A. Lamme et al., 1998; V. A. F. Lamme & Roelfsema, 2000; O’Reilly et al., 2013; Wyatte et al., 2012).

Converging experimental and computational findings support a role for recurrence in challenging recognition. Backward masking has provided influential evidence for such processing by showing that disrupting activity after the initial feedforward response selectively impairs recognition of degraded or ambiguous stimuli (Boehler et al., 2008; Fahrenfort et al., 2007, 2008; V. A. F. Lamme et al., 2002; Ollikka et al., 2026; Wyatte et al., 2012). In an MEG study, (Rajaei et al., 2019) showed that occlusion delayed the emergence of object-category information and that recurrent neural-network models better accounted for recognition under occlusion than purely feedforward models. Related studies have linked difficult recognition to extended neural dynamics and greater recurrent computational depth (Kietzmann et al., 2019; Li et al., 2026; Loke et al., 2022; Spoerer et al., 2020; Tang et al., 2018; Xie et al., 2025). These findings establish the importance of processing beyond the initial response but do not, by themselves, identify the recurrent architecture responsible.

A central unresolved question is whether the additional computation is better explained by local recurrence within visual areas or by long-range top-down feedback between regions. These mechanisms are not mutually exclusive, but they offer different accounts of how incomplete representations are refined. Local recurrent interactions could iteratively integrate visible fragments, resolve boundary ambiguity, and stabilize representations within early and intermediate stages of the ventral hierarchy (Kang et al., 2026; Liao & Poggio, 2020; Linsley et al., 2020; Rajaei et al., 2019; Spoerer et al., 2017). Long-range feedback, in contrast, could convey contextual information or perceptual hypotheses from higher-level regions to earlier processing stages (Bar et al., 2006; Gilbert & Li, 2013) (Kok et al., 2012; T. S. Lee & Mumford, 2003; Morgan et al., 2019).

Evidence for the latter account comes from studies associating demanding recognition with frontal–temporal interactions (Casile et al., 2025; Fyall et al., 2017; Kar et al., 2019; Kar & DiCarlo, 2021), ventrolateral prefrontal-to-temporal communication during occluded-face processing (Noroozi et al., 2024), and feedback within high-level temporal cortex during occluded-body recognition (Bognár et al., 2025). The adaptive-recurrence framework similarly proposes that parieto-frontal networks are recruited when feedforward processing is insufficient (Oyarzo et al., 2025). An important question is therefore whether occlusion-related processing within the ventral stream is accompanied by increased long-range feedback or can be accounted for by local recurrence without a detectable change in interareal communication.

Distinguishing these possibilities requires more than measuring when category information becomes available. A delayed representation may reflect prolonged computation within an otherwise stable circuit rather than a change in the organization of communication between regions. Likewise, sensitivity to backward masking indicates dependence on continued processing but does not uniquely distinguish local recurrence from long-range feedback. Computational comparisons face a related challenge: an advantage for a recurrent model may arise from local interactions, top-down pathways, additional computational depth, or differences in training and architecture (Kang et al., 2026; Liao & Poggio, 2020; Maniquet et al., 2025; Nayebi et al., 2022). Addressing these ambiguities requires convergent evidence from representational dynamics, directed interareal interactions, and controlled comparisons of candidate recurrent architectures.

Here, we combine source-resolved MEG responses from early visual cortex (V1–3), lateral occipital complex (LOC), and inferotemporal/parahippocampal cortex (IT–PHC) with time-resolved decoding (Cichy et al., 2014; Grootswagers et al., 2017; Kaneshiro et al., 2015), backward masking and cross-condition generalization (Xie et al., 2025), and representational Granger causality (Karimi-Rouzbahani et al., 2021; Kietzmann et al., 2019; Rahimi et al., 2023). These analyses assess whether occlusion prolongs category-related processing, whether its late component depends on uninterrupted processing, and whether these temporal effects are accompanied by changes in directed interactions among ventral-stream regions. We further compare three nested readout variants of a trained Hierarchical Top-down Recurrent Network (HTRN): a feedforward variant (HTRN-FF), a locally recurrent variant (HTRN-LR), and a variant with added long-range top-down feedback (HTRN-TD). Because these variants share the same trained backbone, this comparison reduces confounds related to architecture and learned representations, and allows the contributions of local recurrence and top-down feedback to be assessed within a common framework. Together, these analyses test whether local recurrence accounts for the additional behavioral and neural demands of occluded-object recognition and whether the measured and modeled long-range feedback pathways provide additional explanatory value.

## Results

### Occlusion delays category information across the ventral visual pathway

We analyzed source-resolved MEG responses to briefly presented objects from four categories— camel, deer, car, and motorcycle—at three levels of occlusion (0%, 60%, and 80%) (Rajaei et al., 2019). Fourteen participants were included after excluding one participant for excessive recording noise. We examined three anatomically defined regions spanning the ventral visual pathway: V1– 3, lateral occipital cortex (LOC), and inferior temporal–parahippocampal cortex (IT–PHC). Time-resolved pairwise classification quantified category information within each region (Cichy et al., 2014; Grootswagers et al., 2017), with the neural analyses below comparing unoccluded and 60%-occluded objects (Fig. 1A).

**Figure 1.**
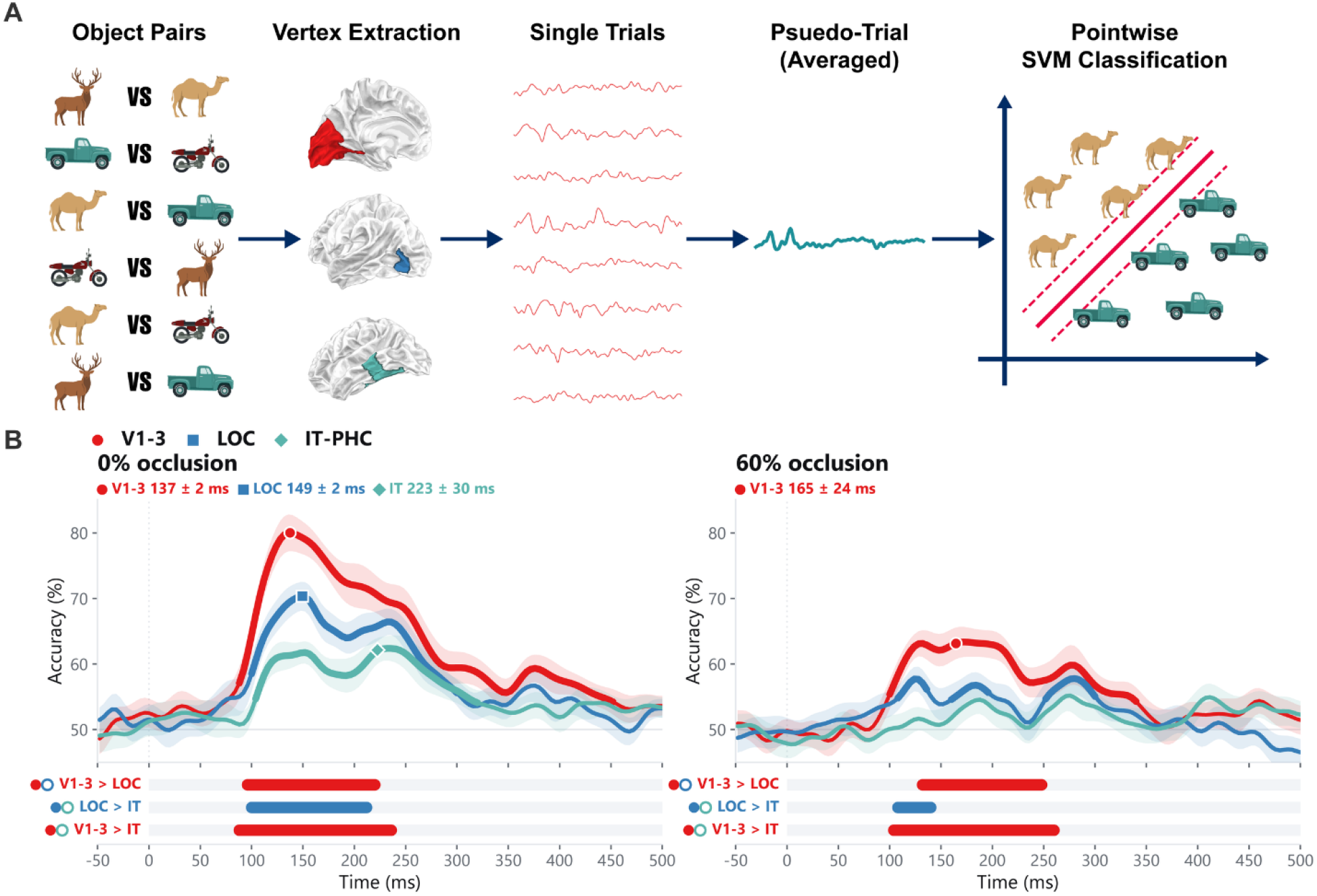
Time-resolved pairwise object-category decoding across ventral visual ROIs. **A)** Schematic of the pairwise decoding analysis. Source-space activity patterns were extracted from vertices within each predefined ROI. Single trials were combined into pseudo-trials, and a linear support-vector machine was trained and tested at each time point to discriminate between pairs of object categories. **B)** Time-resolved object-category decoding accuracy in V1–3, LOC, and IT–PHC under 0% and 60% occlusion (N = 14 participants). Curves indicate the across-participant mean, and shaded regions indicate the standard error of the mean (SEM). Thickened curve segments indicate intervals during which decoding was significantly above the 50% chance level. Labels above the curves indicate peak decoding latency (mean ± SD). Peak latencies are shown only where the peak fell within a significant above-chance cluster; under 60% occlusion this criterion was not met in LOC, whose peak fell between significant clusters, or in IT–PHC, where decoding did not reliably exceed chance. Horizontal bars below each panel indicate intervals with significant between-ROI differences in decoding accuracy (V1–3 versus LOC, LOC versus IT–PHC, and V1–3 versus IT–PHC; cluster-based permutation tests, p < 0.05). Bar colors indicate the ROI with higher decoding accuracy; gray intervals indicate no significant difference.

For unoccluded objects, category information was reliably decoded in all three regions (Fig. 1B). Above-chance decoding extended from 88–452 ms in V1–3, 100–300 ms in LOC, and 104–320 ms in IT–PHC (all *p*_cluster_ < 0.001). Decoding peaked first in V1–3, at 137 ± 2 ms, followed by LOC at 149 ± 2 ms and IT–PHC at 223 ± 30 ms (mean ± SD; all pairwise latency comparisons, *p*≤ 0.001). Decoding accuracy was also highest in V1–3: V1–3 exceeded LOC from 96–220 ms and IT–PHC from 88–236 ms, whereas LOC exceeded IT–PHC from 100–212 ms (all *p*_cluster_ < 0.001).

Occlusion delayed and weakened category decoding, with the clearest temporal shift in V1–3. Decoding onset increased from 87 ± 10 ms to 102 ± 2 ms (*p* < 0.001), and the peak shifted from 137 ± 2 ms to 165 ± 24 ms (*p* = 0.008). Peak accuracy decreased from 80.0% to 63.1%, with decoding remaining near its maximum over approximately 165–210 ms. In LOC, the peak shifted from 149 ± 2 ms to 213 ± 79 ms (*p* = 0.046), and above-chance decoding was restricted to three intervals: 108–136 ms (*p*_cluster_ = 0.035), 160–196 ms (*p*_cluster_ = 0.030), and 252–300 ms (*p*_cluster_ = 0.016). In IT–PHC, decoding did not reliably exceed chance, precluding reliable onset and peak estimates.

Among regions with reliable decoding under occlusion, peak category information remained earlier in V1–3 than in LOC (*p* = 0.048). V1–3 also retained higher decoding accuracy than LOC from 132–248 ms and IT–PHC from 104–260 ms (both *p*_cluster_ < 0.001), whereas LOC exceeded IT–PHC from 108–140 ms (*p*_cluster_ = 0.020). Thus, occlusion delayed category information and reduced its detectability in higher-order regions without reversing the V1–3-to-LOC peak ordering.

These decoding changes were not accompanied by comparable shifts in evoked-response timing (Supplementary Fig. 1). Evoked-response peaks were similar across unoccluded and occluded conditions in V1–3 (79 ± 2 versus 80 ± 2 ms), LOC (85 ± 2 versus 86 ± 2 ms), and IT–PHC (86 ± 2 versus 87 ± 4 ms). LOC and IT–PHC peaks followed V1–3 by approximately 6–7 ms in both conditions (all *p* ≤ 0.001, Wilcoxon signed-rank tests). Occlusion nevertheless reduced evoked-response amplitude in V1–3 from 75–108 ms and in LOC from 99–135 ms (both *p*_cluster_ < 0.05), with no significant difference in IT–PHC.

Together, these analyses distinguish delayed category information from a general delay in regional evoked activity. Reduced response amplitude may contribute to weaker decoding, but the decoding-latency shifts were not mirrored by the timing of the evoked-response peaks.

### Backward masking identifies a late component of category information in V1–3

We next asked whether the delayed category information generalized across backward masking. Backward masking is expected to affect later processing more strongly than the earliest stimulus-driven response (Boehler et al., 2008; Fahrenfort et al., 2007; V. A. F. Lamme et al., 2002; Ollikka et al., 2026; Seijdel et al., 2021; Wyatte et al., 2012). For each region and occlusion level, classifiers were either trained and tested on unmasked trials (“within”) or trained on masked trials and tested on unmasked trials (“cross”; Fig. 2). Because both analyses used unmasked test trials, lower cross-condition accuracy indicates reduced generalization from representations learned under masking to those expressed without a mask (Xie et al., 2025). This comparison measures mask-sensitive category information, although it does not by itself distinguish changes in representational format from reduced information or reliability in the masked training data.

**Figure 2.**
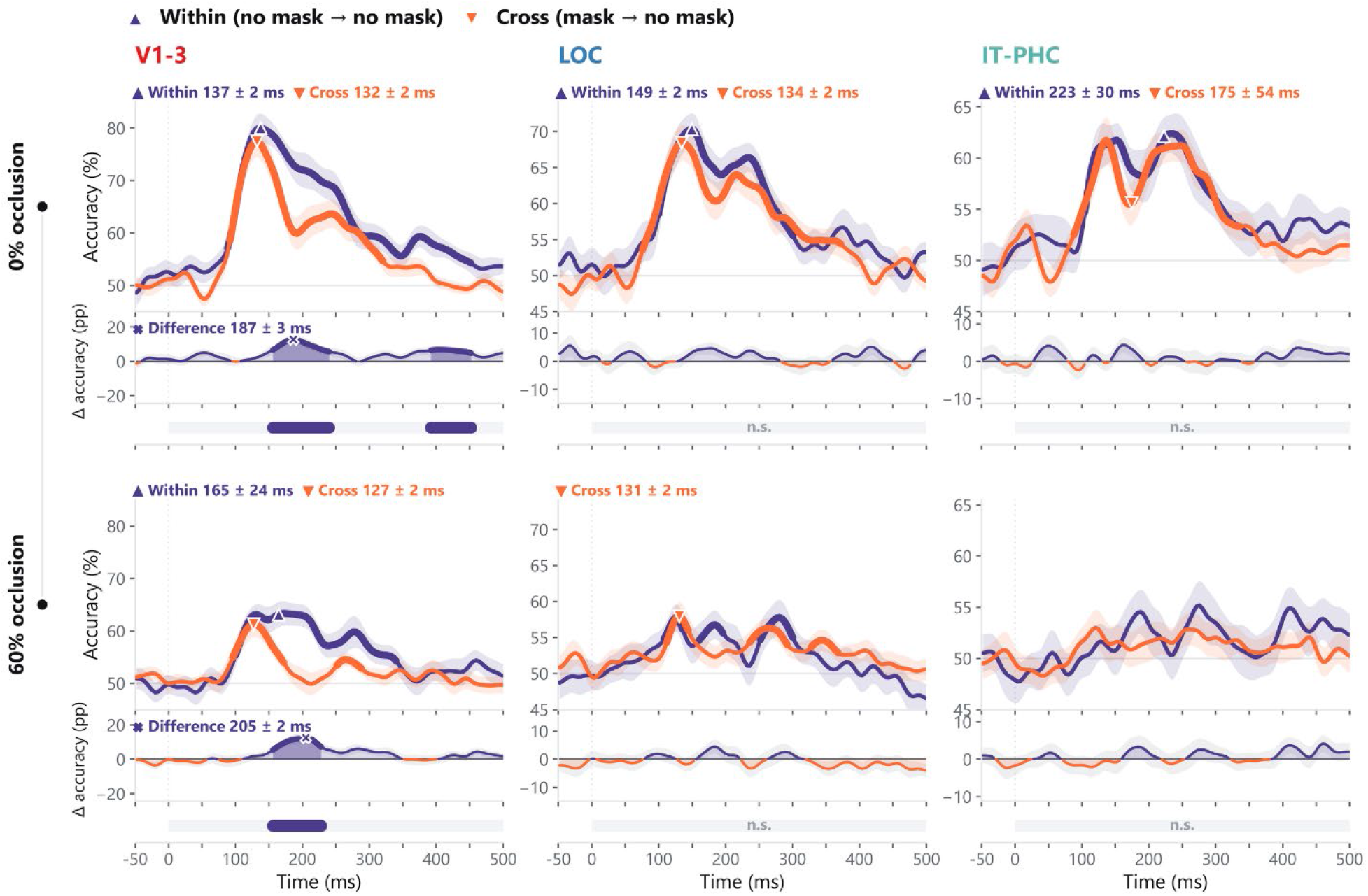
Generalization of object-category decoding across backward masking. For each ROI (V1–3, LOC, and IT–PHC) and occlusion level (0% and 60%), a linear classifier was either trained and tested on unmasked trials (“Within,” no mask → no mask) or trained on masked trials and tested on unmasked trials (“Cross,” mask → no mask). A) Unoccluded objects (0% occlusion). B) 60%-occluded objects. Columns show V1–3, LOC, and IT–PHC (N = 14 participants). Curves indicate the across-participant mean decoding accuracy, and shaded regions indicate the standard error of the mean (SEM). Thickened curve segments indicate intervals of significant above-chance decoding. Labels above the curves indicate peak decoding latencies (mean ± SD), and are shown only where the peak fell within a significant above-chance cluster. Lower panels show the difference between within- and cross-condition decoding accuracy (Δ accuracy, percentage points). In these panels the difference curve is drawn in the within-condition color where within-condition accuracy exceeded cross-condition accuracy and in the cross-condition color where the reverse held; within each significant cluster the area under the curve is shaded and its maximum is marked by a cross, with the adjacent label giving that latency (mean ± SD). Horizontal bars below each panel repeat these significant within–cross difference intervals identified using cluster-based permutation tests (p < 0.05). “n.s.” indicates that no significant cluster was detected.

For unoccluded objects, cross-condition decoding was significant in all three regions: V1–3 from 92–320 ms, LOC from 76–372 ms, and IT–PHC from 88–336 ms (all *p*_cluster_ < 0.001). No within– cross difference was detected during the early 80–150-ms interval, indicating that early object information generalized across masking without a detectable loss in decoding accuracy.

A later difference emerged in V1–3. Within-condition decoding exceeded cross-condition decoding from 156–240 ms (*p*_cluster_ < 0.001), followed by a smaller difference from 392–452 ms (*p*_cluster_ = 0.010). Thus, mask-sensitive category information was present even for unoccluded objects.

Under 60% occlusion, the V1–3 generalization deficit overlapped the delayed decoding peak. Cross-condition decoding peaked at 127 ± 2 ms, earlier than within-condition decoding at 165 ± 24 ms (*p* < 0.001), and within-condition accuracy exceeded cross-condition accuracy from 156– 228 ms (*p*_cluster_ < 0.001). No significant within–cross difference was detected in LOC, and IT–PHC decoding remained unreliable under occlusion.

Backward masking therefore distinguished relatively preserved early generalization from a later generalization deficit in V1–3. This late component occurred in both viewing conditions, but under occlusion it coincided with the delayed decoding peak, linking that delay to information that was less accessible to classifiers trained on masked responses, rather than establishing a processing component unique to occluded recognition.

### Feedback flow is detectable in both conditions, without an occlusion-related change in interareal flow

We then tested whether delayed category information was accompanied by a measurable change in directed interareal interactions. Time-resolved representational Granger causality assessed whether the recent representational history of one region improved prediction of another region’s current representational geometry beyond the target region’s own history (Fig. 3A) (Granger, 1969; Karimi-Rouzbahani et al., 2021; Kietzmann et al., 2019).

**Figure 3.**
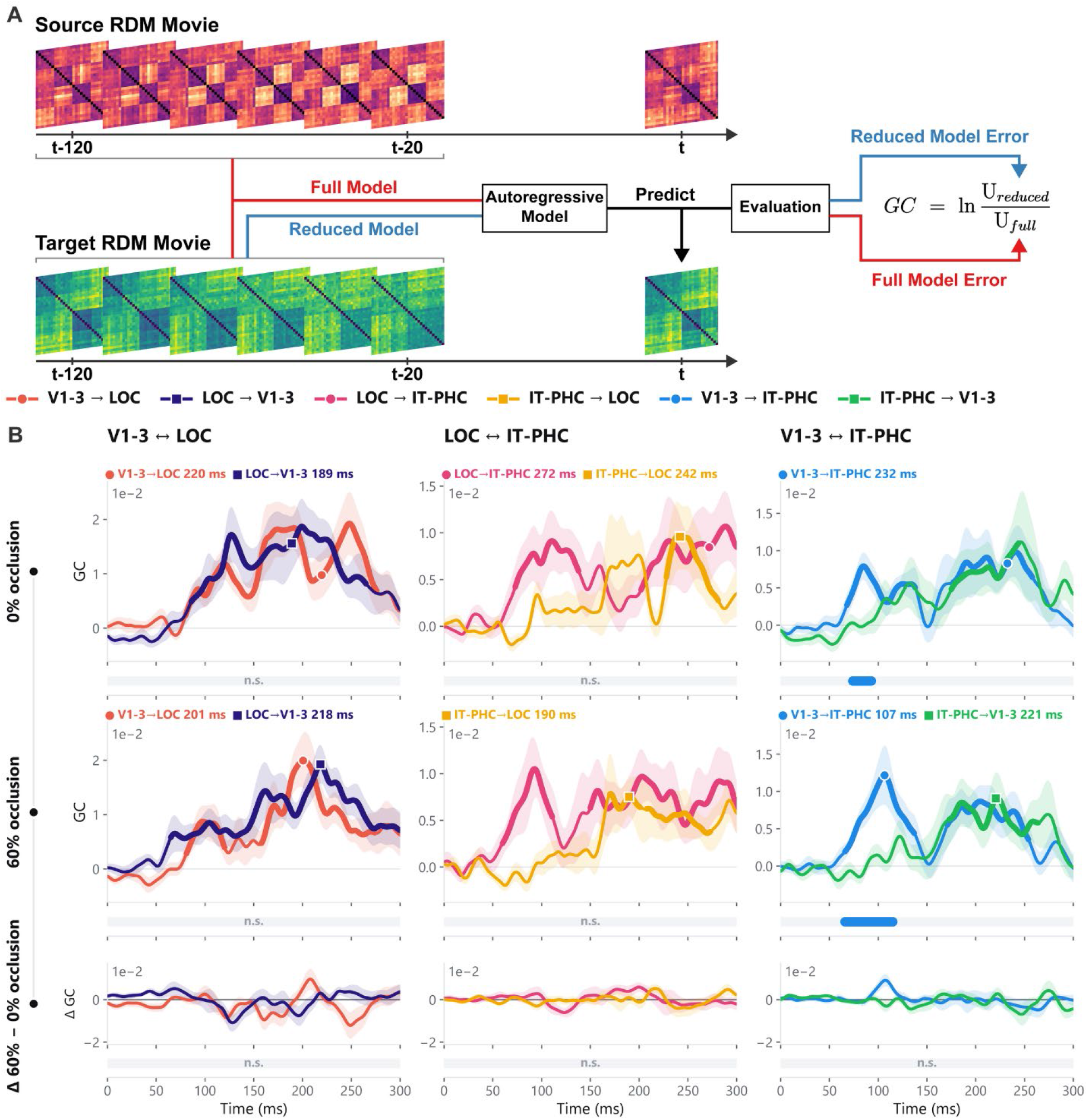
Representational Granger causality between visual ROIs. **A)** Schematic of the time-resolved representational Granger-causality analysis. For each ordered pair of ROIs, the recent history of the source region’s representational geometry was used to predict the target region’s current geometry, after accounting for the target region’s own history. **B)** Grand-average Granger-causality time courses for bottom-up and top-down information flow between V1–3, LOC, and IT–PHC under 0% and 60% occlusion (N = 14 participants). The color key between panels A and B identifies the six directed ROI pairs and applies throughout B. Curves show the mean across participants, and shaded areas indicate the standard error of the mean (SEM). Thickened curve segments indicate significant above-baseline information flow, and labels show peak latencies. Horizontal bars below each panel mark significant differences between the two directions within an ROI pair, identified using cluster-based permutation tests (p < 0.05). Colors indicate the direction with greater information flow; “n.s.” indicates that no significant difference was detected. Bottom panels show the occlusion-related change in each direction of flow (60% minus 0% occlusion), plotted separately for bottom-up and top-down; positive values indicate stronger flow under occlusion.

Between V1–3 and LOC, significant directed interactions were detected in both directions under both viewing conditions (Fig. 3B). Bottom-up flow was significant from 80–276 ms for unoccluded objects (*p*_cluster_ < 0.001) and from 81–120 ms and 158–274 ms under occlusion (*p*_cluster_ = 0.034 and *p*_cluster_ < 0.001, respectively). Top-down flow was significant from 86–265 ms without occlusion and from 65 ms to the end of the analysis window under occlusion (both *p*_cluster_ < 0.001). No significant difference between directions was detected in either condition.

Between LOC and IT–PHC, significant bottom-up flow occurred early, whereas significant top-down flow appeared later. For unoccluded objects, bottom-up flow was detected from 73–138 ms (*p*_cluster_ = 0.004), and top-down flow from 228–281 ms (*p*_cluster_ = 0.012). Under occlusion, the corresponding intervals were 60–111 ms for bottom-up flow (*p*_cluster_ = 0.016) and 170–201 ms and 208–273 ms for top-down flow (*p*_cluster_ = 0.045 and 0.009, respectively).

The V1–3 and IT–PHC pair showed a clear early directional asymmetry. Bottom-up flow was significant from 68–106 ms and 111–138 ms without occlusion (*p*_cluster_ = 0.017 and 0.046) and from 66–137 ms under occlusion (*p*_cluster_ = 0.002). It re-emerged later in both conditions, from 177–258 ms and 171–254 ms, respectively (both *p*_cluster_ = 0.002). Top-down flow was detected during the later response, from 175–244 ms without occlusion (*p*_cluster_ = 0.008) and 173–264 ms under occlusion (*p*_cluster_ = 0.004). Bottom-up flow exceeded top-down flow early in both conditions: 74–93 ms without occlusion (*p*_cluster_ = 0.049) and 66–115 ms under occlusion (*p*_cluster_ = 0.005).

Despite these detectable interactions, direct comparisons revealed no significant occlusion-related difference in either bottom-up or top-down flow for any ROI pair. The per-direction occlusion contrasts likewise provided no evidence for a selective increase in top-down flow under occlusion (Fig. 3B, bottom). Backward masking produced localized reductions in late flow but no consistent change across interareal connections (Supplementary Figs. 2–4).

These findings distinguish the presence of feedback from an occlusion-related change in feedback. Top-down interactions were detectable in both conditions; the null between-condition result therefore does not imply that feedback was absent during occluded recognition. Rather, the delayed category information occurred without a detectable change in the directed interareal interactions captured by this analysis.

### Local recurrence recovers recognition performance as occlusion increases

To test which model computations could account for recognition under occlusion, we compared three nested variants of a Hierarchical Top-down Recurrent Network: a feedforward variant (HTRN-FF), a locally recurrent variant (HTRN-LR), and a variant combining local recurrence with long-range top-down feedback (HTRN-TD; Fig. 4A, 4B). Stage 0 provided the feedforward readout; subsequent stages progressively incorporated recurrent computation.

**Figure 4.**
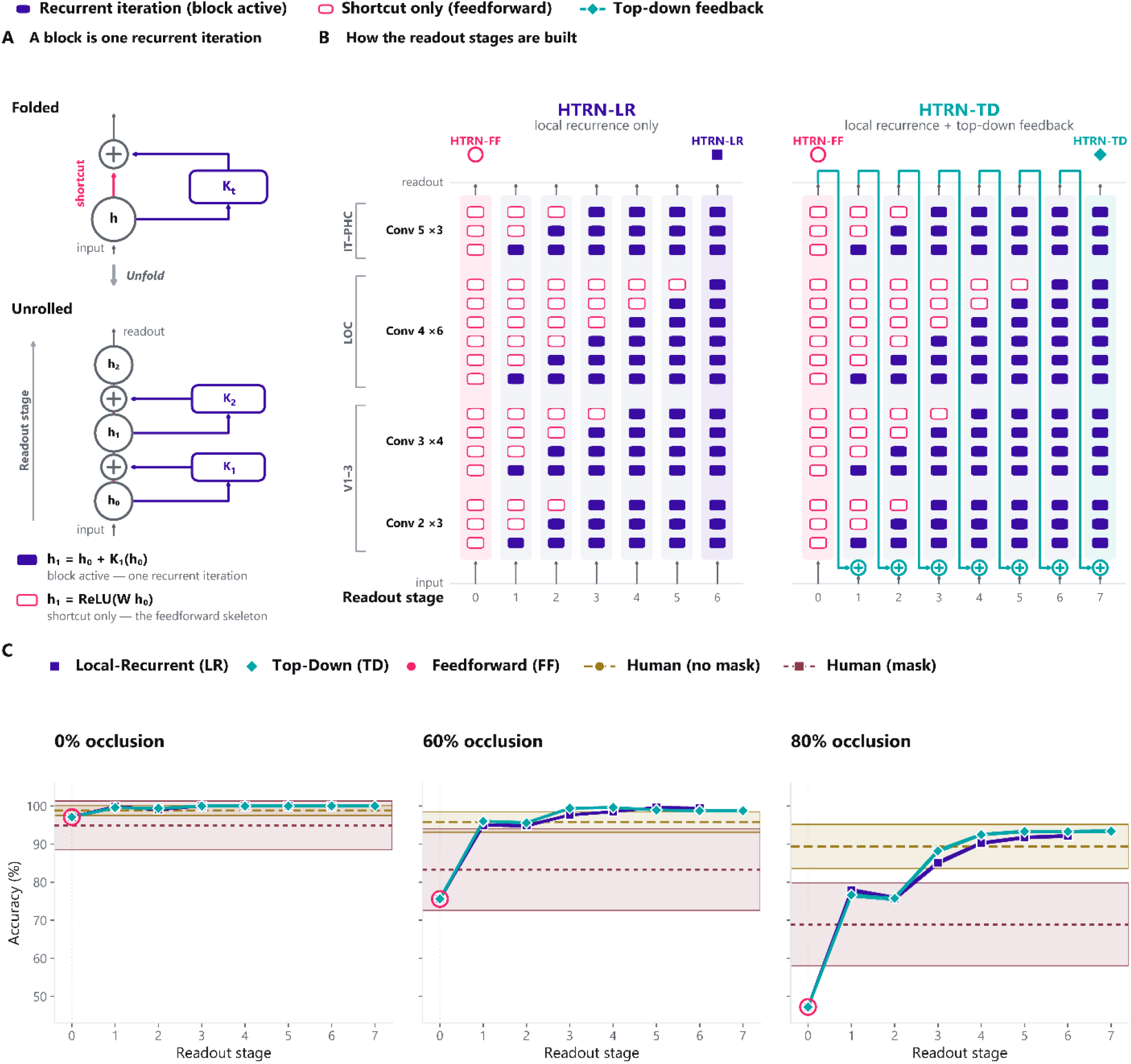
HTRN readout stages and recognition accuracy under occlusion. **A)** A residual bottleneck block interpreted as one recurrent iteration. In the folded representation, the block applies a recurrent transformation to its input and combines the result with the shortcut pathway. Unrolling the block exposes successive computational states. An active bottleneck block contributes one recurrent iteration (filled purple), whereas the projection shortcut preserves the feedforward pathway without adding recurrent computation (open magenta). **B)** Construction of the HTRN readout stages across the bottleneck blocks of the ResNet-50 backbone. Stage 0 corresponds to the feedforward readout. Later stages progressively add local recurrent computation. HTRN-LR contains local recurrence only, whereas HTRN-TD incorporates additional long-range top-down feedback during the recurrent processing sequence. **C)** Four-category classification accuracy across readout stages under 0%, 60%, and 80% occlusion, obtained from a multiclass linear classifier trained on globally average-pooled features from each readout stage (8-fold stratified cross-validation, repeated 20 times). Curves show the mean accuracy across folds and repetitions; shaded regions indicate the SEM across repetitions and are often narrower than the line width. Dashed lines show human performance in the no-mask and mask conditions, with surrounding bands spanning the error bars reported for those estimates (behavioral accuracies adopted from Rajaei et al., 2019). Chance performance was 25%.

For unoccluded objects, feedforward recognition was already near ceiling, leaving limited scope for further improvement (Fig. 4C). Occlusion exposed a substantial benefit from additional computation. At 60% occlusion, accuracy increased from 75.6% at the feedforward stage to approximately 96% after the first recurrent stage and approached ceiling at later stages. At 80% occlusion, accuracy increased from 47.2% to approximately 77% after one recurrent stage and continued to improve, reaching 92.1% for HTRN-LR. The readout sequence thus showed that local recurrence recovered much of the recognition performance lost as visible object information decreased.

Adding long-range top-down feedback yielded similar performance to local recurrence alone. Final accuracies were 99.5% for HTRN-LR and 98.8% for HTRN-TD at 60% occlusion, and 92.1% and 93.4%, respectively, at 80% occlusion. The implemented top-down pathway therefore produced no consistent additional accuracy gain across occlusion levels.

Occlusion robustness differed across the other architectures tested (Supplementary Fig. 5). Although all models performed at or near ceiling for unoccluded objects (97.1–100%), their performance diverged at 80% occlusion. BLT-FF, BLT-LR, and BLT-TD achieved 63.4%, 70.0%, and 64.2% accuracy, respectively, whereas the feedforward and recurrent CORnet variants achieved 65.8% and 50.9%. In contrast, the recurrent HTRN variants remained numerically close to unmasked human performance and above masked human performance at both occlusion levels.

These comparisons identified HTRN-LR and HTRN-TD as the most occlusion-robust models among those tested. Their nested architecture also allowed us to ask whether the computations supporting this performance captured the temporal evolution of human neural representations.

#### Local recurrence captures late, mask-sensitive representations in early visual cortex

We next asked whether the computations supporting occlusion-robust recognition also captured the temporal evolution of neural category representations. We compared model and MEG representational geometries over time, matching HTRN Conv2–3 representations to V1–3, Conv4 to LOC, and Conv5 to IT–PHC (Fig. 5A). The nested model variants allowed us to assess the contribution of local recurrence beyond feedforward computation and the additional benefit of the implemented long-range top-down pathway.

**Figure 5.**
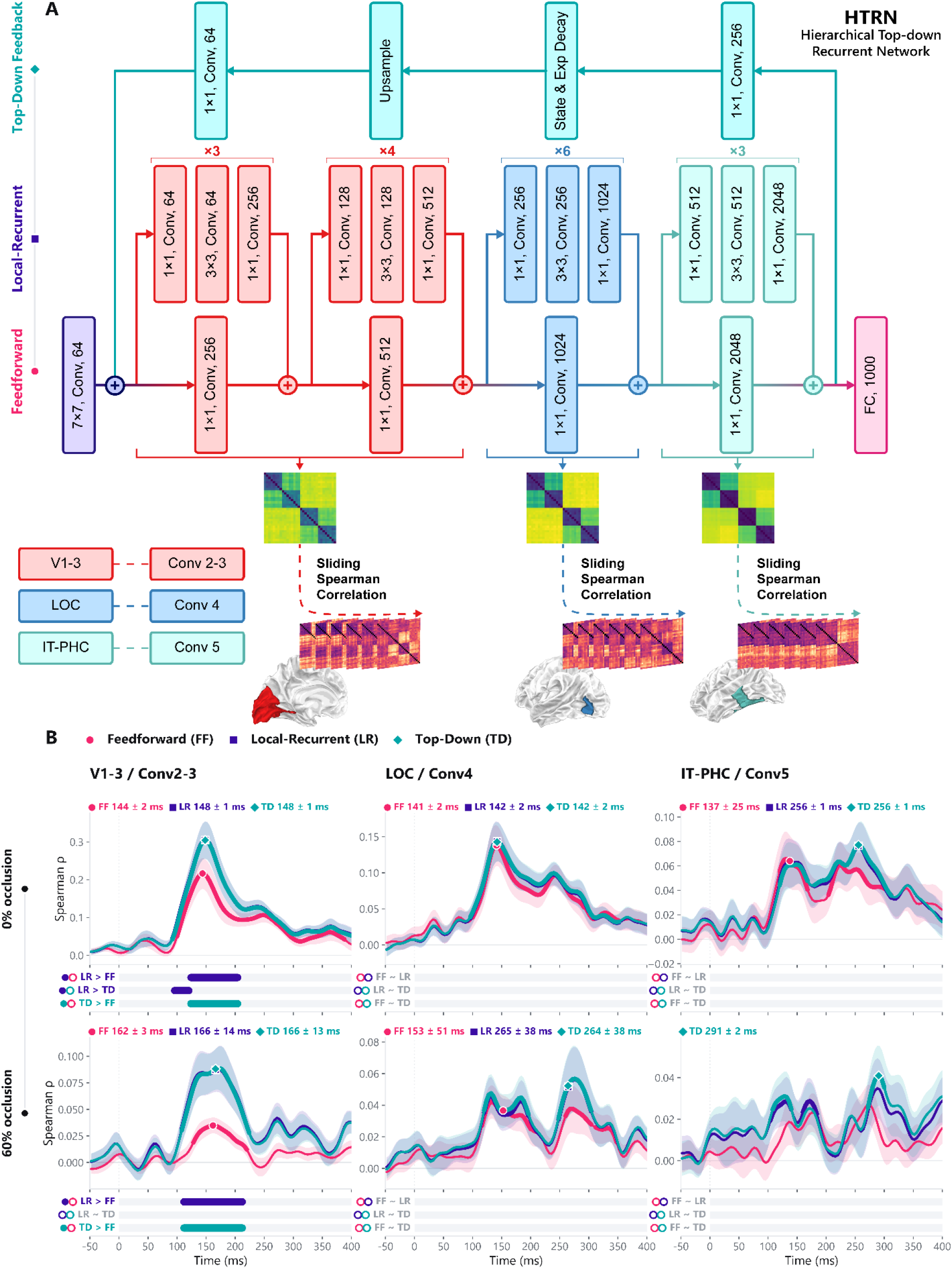
Model–brain representational similarity across feedforward and recurrent computations. **A)** Architecture of the Hierarchical Top-down Recurrent Network (HTRN) and model–brain comparison. The trained network was evaluated as three nested variants: a feedforward model (HTRN-FF), a model with local recurrent computation (HTRN-LR), and a model combining local recurrence with long-range top-down feedback (HTRN-TD). Representations from Conv2–3, Conv4, and Conv5 were compared with source-localized MEG representations from V1–3, LOC, and IT–PHC, respectively. At each neural time point, model and neural representational dissimilarity matrices were compared using a sliding Spearman correlation. **B)** Time-resolved model–brain correspondence under 0% and 60% occlusion (N = 14 participants). Curves show mean Spearman correlations across participants, and shaded regions indicate the SEM. Labels report peak latencies (mean ± SD). Thickened curve segments indicate significant positive model–brain correlations. Horizontal bars below each panel mark significant pairwise differences among HTRN-FF, HTRN-LR, and HTRN-TD, identified using cluster-based permutation tests (pcluster < 0.05).

For unoccluded objects, all three HTRN variants correlated significantly with V1–3 representations from approximately 100 ms onward (*p*_cluster_ < 0.001). Both recurrent variants nevertheless showed stronger correspondence than HTRN-FF from 124–204 ms (both *p*_cluster_ < 0.001; Fig. 5B). No significant differences among variants were detected in LOC or IT–PHC. Thus, recurrent computation improved neural correspondence even when feedforward recognition performance was already near ceiling.

Under 60% occlusion, HTRN-FF showed weak correspondence with V1–3, whereas both recurrent variants exceeded it from 112–212 ms (both *p*cluster < 0.001). This interval overlapped the delayed category-decoding peak and the late masking-related generalization deficit. HTRN-TD did not significantly outperform HTRN-LR, and no reliable differences among variants were detected in LOC or IT–PHC. Local recurrence therefore captured the observed improvement in model–brain correspondence in the late V1–3 response, without a detectable additional benefit from the implemented top-down pathway. Masking reduced correspondence with both recurrent variants from 140–196 ms for unoccluded objects and from 176–212 ms under occlusion (all *p*cluster < 0.05; Supplementary Figs. 6–7). No significant masking effects were detected in LOC or IT–PHC. The recurrent-model advantage thus converged spatially and temporally with mask-sensitive category information, including the interval surrounding the delayed decoding peak under occlusion.

Comparisons with additional architectures showed that the benefit of recurrence depended on its implementation (Supplementary Figs. 8–11). In BLT-VS, local recurrence improved model–brain correspondence in LOC, but adding top-down feedback provided no further significant advantage. In CORnet, recurrence improved neither recognition performance nor neural correspondence. Across the tested model families, no reliable representational benefit of additional top-down processing was detected.

Although adding the implemented top-down pathway yielded no consistent benefit over local recurrence in the occlusion task, the full HTRN-TD model improved recognition performance on the broader ImageNet-1k validation set relative to its original pretrained ResNet-50 backbone. Top-1 accuracy increased from 77.03% to 78.86% (*p* < 0.001; bootstrap 95% CI of the difference, 1.57–2.10 percentage points), and top-5 accuracy increased from 93.76% to 94.36% (95% CI of the difference, 0.44–0.75 percentage points). Thus, the absence of an additional top-down benefit under occlusion occurred in a model that nevertheless improved large-scale recognition performance over its feedforward backbone.

Together, the results connect delayed category information under occlusion to a late, mask-sensitive V1–3 representation that was better captured by local-recurrent than feedforward computations. Feedback interactions were detectable in both viewing conditions, but occlusion produced no detectable change in the measured interareal flow. Within the tested HTRN architecture, local recurrence recovered occlusion-robust recognition and improved late neural correspondence, without a consistent additional benefit from the implemented long-range top-down pathway.

## Discussion

Recognizing occluded objects requires computation beyond the initial feedforward sweep, but the form of that additional computation remains unclear. Our findings favor prolonged local recurrent processing within the ventral visual hierarchy over an additional contribution from the long-range top-down pathway tested here. Occlusion delayed and prolonged category-related processing without measurably altering the temporal ordering or directed interactions among the ventral-stream regions examined. In the computational comparisons, local recurrence improved recognition under occlusion and correspondence with late neural dynamics, whereas adding the implemented long-range top-down pathway provided no consistent additional explanatory benefit. Together, these findings constrain both the temporal characteristics and the possible architecture of the recurrent computations supporting occluded-object recognition.

### Prolonged processing without detectable reorganization of ventral-stream interactions

A key implication of these results is a dissociation between the duration of visual processing and the organization of interareal communication. Under occlusion, category information emerged later and persisted longer, but this temporal extension was not accompanied by a detectable change in the relative progression of responses across V1–3, LOC, and IT–PHC or in their directed interactions. Delayed recognition therefore need not imply recruitment of a distinct feedback route. Instead, the present findings are consistent with recurrent computation increasing effective processing depth while preserving the broad organization of ventral-stream processing (Kubilius et al., 2019; Liao & Poggio, 2020; Nayebi et al., 2022; Spoerer et al., 2020).

This distinction is important because prolonged neural responses alone do not identify the mechanism responsible for additional processing. The extended dynamics observed under occlusion could reflect repeated computation within an existing network rather than a measurable redistribution of communication between regions. Our results support this account within the spatial and representational scope of the present analyses, although they do not establish that interareal communication remains unchanged in all respects.

### Backward masking constrains the interpretation of late category information

Backward masking provides complementary evidence about the late occlusion-sensitive component. Early category information generalized between masked and unmasked trials, whereas the later component was selectively disrupted by masking. This pattern is consistent with a late stage of processing that depends on continued processing after the initial representation has emerged. It also suggests that the effects of masking are temporally selective rather than a uniform reduction of category information throughout the response.

The overlap between this mask-sensitive interval and the period in which recurrent models outperformed the feedforward model further links the late component to iterative computation. However, masking alone cannot distinguish local recurrence from long-range feedback or identify the representational operation being disrupted. Nor does selective disruption of late information, by itself, establish whether recurrence transforms the representation, stabilizes it, or sustains its decodability. The mechanistic interpretation therefore rests on the convergence of masking, directed-connectivity, and model–brain comparisons rather than on masking as an isolated assay of recurrence.

### Local recurrence and the resolution of occlusion-related ambiguity

The recurrent-model advantage was strongest in V1–3, highlighting early visual cortex as a potential contributor to resolving occlusion-related ambiguity. This regional pattern does not establish where recurrent processing begins, but it suggests that its contribution is not restricted to high-level object representations. Occlusion creates ambiguity about whether contours and image fragments belong to the object or the occluder (Johnson & Olshausen, 2005; Namima & Pasupathy, 2021; Rensink & Enns, 1998). Local recurrent interactions may help resolve this ambiguity by integrating visible fragments, incorporating contextual information, stabilizing object boundaries, or suppressing occluder-related features (Choi et al., 2018; Kosai et al., 2014; Tang et al., 2018).

These computations could support contour completion, iterative segmentation, or “explaining away,” whereby an occluder representation accounts for missing or disrupted object features and facilitates integration of the remaining evidence (Ernst et al., 2021; Kang et al., 2026; Kok et al., 2012; T. S. Lee & Mumford, 2003; Namima & Pasupathy, 2021). Such interpretations are consistent with accounts in which fragmented input is progressively transformed into integrated object- or shape-based representations (Ban et al., 2013; Erlikhman & Caplovitz, 2017; Hegdé et al., 2008; Rauschenberger et al., 2006; Seijdel et al., 2021; Tang et al., 2014), but the present data do not distinguish among these recurrent operations. In particular, a better match between local-recurrent models and neural representations does not uniquely identify the algorithm implemented by biological circuits. Targeted manipulations of occluder geometry, contour continuity, and fragment diagnosticity could help separate contributions from boundary assignment, fragment integration, and representational stabilization.

### Model comparisons and the scope of the top-down conclusion

The nested modeling framework strengthens the comparison between candidate recurrent architectures. HTRN-FF, HTRN-LR, and HTRN-TD shared the same trained backbone and differed in the recurrent computations they implemented, reducing confounds associated with comparing independently trained architectures. Within this common framework, local recurrence improved recognition under occlusion and increased correspondence with human ventral-stream dynamics during the late, mask-sensitive interval, particularly in V1–3. Adding the implemented top-down pathway did not provide a consistent further benefit.

This lack of additional benefit is not readily attributable to an entirely nonfunctional feedback mechanism: the same pathway improved ImageNet-1k validation performance over the pretrained ResNet-50 backbone (Calhas & Oliveira, 2025). That improvement demonstrates computational utility in another evaluation setting, although it does not establish that the pathway captures the feedback computations relevant to human occluded-object recognition. The model comparison therefore supports local recurrence as a sufficient mechanism within the tested framework, rather than ruling out alternative implementations of top-down processing.

The neural analyses place a similarly specific constraint on the role of feedback. We found no occlusion-related increase in measured directed feedback within the ventral pathway, but this should not be interpreted as evidence that feedback is absent or unimportant. Representational Granger causality may miss feedback that modulates gain, precision, attention, or features not captured by the category-related representations analyzed here (Groen et al., 2018; Kok et al., 2012, 2016; Morgan et al., 2019). Moreover, our analyses focused on selected ventral-stream regions and did not directly test contributions from frontal or parietal cortex (Bar et al., 2006; Carricarte et al., 2025; Oyarzo et al., 2025).

Long-range feedback may contribute more strongly under greater degradation or near-threshold recognition demands (Ben-Yosef et al., 2018; Bognár et al., 2025; Casile et al., 2025; Fyall et al., 2017; Kar & DiCarlo, 2021; Noroozi et al., 2024; Ullman et al., 2016). The more precise conclusion is therefore that, under the present challenging but behaviorally manageable conditions, robust recognition occurred without a detectable increase in the measured long-range feedback interactions, and the implemented top-down pathway did not improve the account of the neural data.

### Conclusions

Occlusion delayed and prolonged category-related processing without measurably altering the broad organization of ventral-stream interactions. The late component was selectively vulnerable to masking and was better captured by local-recurrent than feedforward model dynamics, with no consistent further benefit from the implemented long-range top-down pathway. Together, these findings favor prolonged local recurrent processing as an account of the additional computation supporting recognition in the present task. This conclusion is specific to the regions, signals, architectures, and task demands examined and does not exclude contributions from broader top-down interactions.

## Methods

### Objects image set

This study used the previously established MEG-Occlusion dataset (Rajaei et al., 2019; Rajaei & Khaligh-Razavi, 2019). The stimulus set comprised images from four object categories: camel, deer, car, and motorcycle. Each category was presented under systematically manipulated levels of visual occlusion.

Occluded stimuli were generated by iteratively placing artificial occluders over both the foreground and background regions of the target objects. This procedure yielded three predefined occlusion levels: 0% (unoccluded), 60%, and 80%. To control for potential effects of the occluders themselves, the unoccluded condition also contained identical black disks; however, in this condition the disks were positioned so that they did not cover the target objects.

The full design comprised 24 experimental conditions (4 object categories × 3 occlusion levels × 2 masking conditions). Each combination of category and occlusion level was represented by a unique set of 64 exemplars, varying in object position and in the placement of the occluders. To reduce the contribution of low-level visual features, target object locations were randomly jittered around the image center, and all images were normalized for size and contrast. On unmasked trials only the target image was shown; on masked trials the target image was immediately followed by a dynamic backward mask, following the masking manipulation of the original MEG-Occlusion paradigm (Rajaei et al., 2019).

All three occlusion levels were presented during MEG acquisition. The MEG analyses reported here contrast the 0% and 60% conditions. The 80% condition is retained in the model evaluations, where it provides the most demanding test of recurrent computation.

### Participants and experimental design

Fifteen healthy adults participated in the experiment; one participant was excluded because of excessive noise in the MEG recording, yielding a final sample of 14 participants (age range: 23– 38 years; 7 females; all right-handed). To investigate the spatiotemporal dynamics of object recognition under occlusion, participants completed a rapid visual presentation task. Each trial began with a central fixation cross presented for 1 s, followed by an object image displayed for 34 ms. On masked trials, the object image was followed, after a 17-ms inter-stimulus interval, by a dynamic backward mask presented for 102 ms (a sequence of six texture-synthesized images, each shown for 17 ms), giving a target-to-mask stimulus-onset asynchrony of 51 ms. To maintain task engagement and encourage attention to object category, a question trial was presented after every 1–3 object trials, in which participants were asked to indicate whether the object shown on the immediately preceding trial belonged to the animal or non-animal category.

### MEG data acquisition and preprocessing

MEG data were recorded using a 306-sensor Elekta Neuromag system. Signals were sampled at 1000 Hz and acquired with an online hardware band-pass filter of 0.03–330 Hz. Data preprocessing was performed using MaxFilter (Taulu & Simola, 2006), MNE-Python (Gramfort et al., 2013), and the Brainstorm toolbox (Tadel et al., 2011).

Environmental noise and head-movement artifacts were corrected using spatiotemporal signal space separation (Taulu & Simola, 2006). The data were subsequently band-pass filtered between 1 Hz and 100 Hz. To further remove physiological and non-physiological artifacts, independent component analysis (ICA) was applied using the extended infomax algorithm (T.-W. Lee et al., 1999), and artifactual components were identified and removed manually.

The cleaned data were then epoched from −200 ms to 1000 ms relative to stimulus onset, and baseline correction was applied using the −200 to 0 ms prestimulus interval. Epochs containing residual artifacts were rejected using peak-to-peak amplitude thresholds of 4000 fT/cm for gradiometers and 4 pT for magnetometers. The design specified 64 trials per condition, giving 1,536 trials per participant across the 24 conditions. Across participants, 3.5 ± 4.6% of epochs were rejected (mean ± SD), leaving 1,448 ± 86 trials per participant and 60 ± 4 trials per condition. The data were then temporally smoothed using a 20 Hz low-pass filter.

### Source estimation

Cortical source estimates were reconstructed from sensor-level MEG data using L2 minimum-norm estimation (Hämäläinen & Ilmoniemi, 1994) implemented in MNE-Python (Gramfort et al., 2013). Individual structural MRIs were not available for this dataset, so all analyses used the standard fsaverage template brain. A surface-based source space was generated on fsaverage at oct6 resolution (4098 vertices per hemisphere, approximately 4.9 mm average spacing). The forward solution, which defines the mapping between source dipoles and MEG sensors, was computed using a three-layer boundary element model (BEM) to provide an anatomically realistic approximation of head geometry.

Subject-specific inverse operators were computed from the forward model and a noise covariance matrix estimated from the prestimulus baseline interval (−200 to 0 ms) of all epochs using a shrunk covariance estimator (Engemann & Gramfort, 2015). Inverse operators were constructed with a loose orientation constraint of 0.2 and depth weighting of 0.8. Source estimates were obtained with the L2 minimum-norm solution (MNE), using a regularization parameter of λ² = 1/9, corresponding to an assumed signal-to-noise ratio of 3, and retaining the dipole component normal to the cortical surface. These inverse operators were applied to the preprocessed sensor-level data to obtain source-space activity estimates.

### Regions of interest

To examine representational dynamics across the visual processing hierarchy, we restricted our analyses to three anatomically defined regions of interest (Kietzmann et al., 2019). Cortical parcels were defined according to the Human Connectome Project multimodal parcellation (HCPMMP1) atlas (Glasser et al., 2016) and projected onto the fsaverage cortical surface. Neighboring atlas labels were then merged to form three broader functional ROIs corresponding to early, intermediate, and high-level visual cortex.

The early visual cortex (V1–3) ROI comprised labels V1, V2, and V3. The lateral occipital complex (LOC) ROI was defined by combining LO1, LO2, LO3, and V4t. The inferotemporal/parahippocampal cortex (IT–PHC) ROI was formed by merging higher-level ventral temporal and parahippocampal labels, including PHA1, PHA2, PHA3, VMV2, VMV3, VVC, FFC, TE1p, and TE2p.

These ROIs were intentionally defined as relatively large and spatially distinct regions in order to maximize the signal-to-noise ratio (SNR) while minimizing potential spatial leakage between neighboring cortical areas, a known limitation of source-reconstructed MEG connectivity analyses (Farahibozorg et al., 2018; Hauk et al., 2022).

### ROI-based evoked-response extraction

To characterize the temporal dynamics of regional responses across occlusion levels, ROI time courses were extracted from the source-space data. For each participant and occlusion condition, preprocessed sensor-level epochs were averaged across trials to obtain condition-specific evoked responses, and the inverse operator was then applied to these sensor-level averages to reconstruct source-level activity. No further baseline correction was applied in source space; the epoch-level baseline correction described above was the only one used.

To derive a single representative time course for each ROI (V1–3, LOC, and IT–PHC), source activity was pooled across vertices. Specifically, vertex-wise time courses were extracted and averaged using the mean flip method implemented in MNE-Python (Gramfort et al., 2013). This procedure aligns the signs of the source signals according to local cortical surface orientation prior to averaging, thereby reducing signal cancellation arising from variability in dipole orientations across the folded cortical geometry.

### Multivariate pattern analysis

To evaluate the temporal dynamics of object information across ROIs and levels of occlusion, we performed time-resolved pairwise decoding on source-estimated activity within each ROI (Fig. 1A). To improve signal-to-noise ratio, single trials were randomly assigned to eight bins, and trials within each bin were averaged to form pseudo-trials (Grootswagers et al., 2017; Guggenmos et al., 2018; Isik et al., 2014). This procedure yielded eight pseudo-trials per object category for each pairwise classification analysis.

At each time point, a linear Support Vector Machine (SVM) (Cortes & Vapnik, 1995) was trained and tested using the pattern of activation across all vertices in the ROI as the feature vector. Decoding was performed separately for all six pairwise combinations of the four object categories. Classifier performance was evaluated using an 8-fold stratified cross-validation, yielding a time-resolved measure of pairwise decoding accuracy.

To reduce dependence on random trial assignment during pseudo-trial generation, the full binning and cross-validation procedure was repeated 100 times. Decoding accuracies were then averaged across iterations and across the six object-category pairs. This resulted in a single continuous time course of object-information decoding for each participant, ROI, and occlusion condition.

To evaluate whether object information generalized across the masking manipulation, we performed an analogous cross-condition decoding analysis (Xie et al., 2025) using the same pseudo-trial and shuffling procedure. For each ROI and occlusion level, classifiers were trained on pseudo-trials from masked trials and tested on pseudo-trials from unmasked trials of the same category pair (train-mask/test-no-mask, ‘Cross’), and compared against a within-condition control trained and tested entirely on unmasked trials (‘Within’). The Within − Cross difference in decoding accuracy was computed at each time point to quantify the extent to which object representations depended on the masking manipulation.

### Representational similarity analysis

#### Brain RDM extraction

To characterize the evolving representational geometries within each ROI, we constructed time-resolved representational dissimilarity matrices (RDMs) (Kriegeskorte et al., 2008). At each time point, the multivariate spatial activation pattern across vertices within the ROI was extracted, preserving the spatial structure of the response. Pairwise dissimilarity between condition-specific activation patterns was then computed using Pearson correlation distance (1 − r) (Guggenmos et al., 2018; Kriegeskorte et al., 2008). Repeating this procedure across time yielded a dynamic sequence of RDMs for each ROI, capturing the temporal evolution of representational structure.

#### Model RDM extraction

Model RDMs were built from the images each participant actually viewed. For every participant, occlusion level and object category, the exact stimulus sequence presented during the MEG session was replayed through the network, so that model and neural patterns were aligned trial by trial. Images were resized to 224 × 224 pixels and normalized with the standard ImageNet channel statistics. Unit activations were read out from the network stage assigned to each ROI (see Computational modeling), flattened into a single feature vector per image, and used without further dimensionality reduction.

Model and neural RDMs were then constructed by an identical procedure. Within each condition, the trials of each object category were randomly partitioned into eight groups and averaged, yielding eight sub-averaged patterns per category and therefore a 32 × 32 RDM (496 unique pairs); for the MEG data this was repeated at every time point. Dissimilarity was defined as Pearson correlation distance (1 − r), as for the brain RDMs above. Because the same random partition was applied to the model and to the MEG patterns, corresponding cells of the two RDMs were computed from the same trials.

#### Model–brain representational similarity

Model–brain correspondence was quantified as the Spearman rank correlation between the vectorized off-diagonal elements (496 unique pairs) of the model RDM and those of the MEG RDM at each time point, giving one correlation time course per participant, ROI, model variant and occlusion level. Before RDM construction, each MEG source time course was scaled by its own prestimulus standard deviation, so that vertices contributed to the pattern in proportion to their stimulus-evoked rather than their absolute amplitude.

To reduce dependence on the random assignment of trials to sub-averaging groups, the complete procedure was repeated 100 times with independent partitions, and the resulting correlation time courses were averaged. Group-level statistics were computed across the 14 participants with the cluster-based permutation procedure described below, both against zero and for the pairwise contrasts between HTRN-FF, HTRN-LR, and HTRN-TD. The representational similarity analysis was restricted to unmasked trials and to the 0% and 60% occlusion levels, matching the decoding analyses; the corresponding masked-trial analyses are reported in the supplementary material.

#### Representational Granger causality

To examine the directional flow of information between visual regions, we applied representational Granger causality analysis (Granger, 1969; Kietzmann et al., 2019) to the ROI-specific RDM time series.

At each time point, the current RDM of the target ROI was modeled using linear regression. Causal influence was quantified by comparing two models: a reduced model, which included only the past time window of the target RDM, and a full model, which included the past time windows of both the target and source RDMs. The past time window was defined as −120 to −20 ms relative to the time point being predicted. Granger causality was defined as:

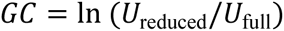

where *U* denotes the unexplained variance (prediction error) of the corresponding models (Fig. 3A). Positive values therefore indicate that incorporating the past representational state of the source region improves prediction of the target region’s current representational geometry.

### Computational modeling

To investigate the computational basis of object recognition under occlusion, we constructed a single model family, the Hierarchical Top-down Recurrent Network (HTRN), from which three nested processing variants were derived: a purely feedforward variant (HTRN-FF), a variant with local recurrent connections (HTRN-LR), and a variant additionally equipped with a long-range top-down feedback pathway (HTRN-TD). All three variants are readout stages of one trained network rather than separately trained models, so that differences between them reflect the presence or absence of a given computational mechanism rather than differences in training, architecture, or capacity.

The backbone of the HTRN follows the conceptual framework established by Rajaei et al. (2019), in which a residual neural network (He et al., 2015) is interpreted as a hierarchical architecture whose residual blocks can be reformulated as local recurrent computation (Liao & Poggio, 2020), thereby providing a model of recurrent processing in the ventral visual stream. HTRN-LR implements this hierarchical recurrent formulation on a pretrained ResNet-50: the residual architecture is treated as implementing iterative local computations across hierarchical stages rather than as a purely feedforward cascade. HTRN-FF approximates the initial feedforward sweep and corresponds to readout stage 0 of the same network, restricting the model to its non-iterative pathway and thereby removing the deeper residual computations while preserving the overall hierarchical structure.

HTRN-TD extends HTRN-LR with a long-range top-down feedback pathway inspired by the Deep Feedback Model (Calhas & Oliveira, 2025), a class of recurrent architectures in which a global, high-level state is iteratively fed back to early processing stages over time. In the HTRN, the network’s final-layer output is compressed into a lower-dimensional feedback state, softmax-normalized, and projected back into the entry layer of the same ResNet-50 backbone used by HTRN-FF and HTRN-LR, where it is additively combined with the bottom-up input; the resulting state is stabilized across iterations by an exponential-decay mechanism that ensures convergence to an equilibrium representation. Relative to the publicly released DFM implementation, we modified the feedback pathway to be additive rather than channel-concatenated, keeping the backbone dimensionally identical to a standard ResNet-50 and compatible with pretrained ImageNet weights, and reduced the feedback state to a compact 256-dimensional bottleneck decoupled from the average pooling features, lowering the memory and compute cost of unrolling the feedback loop.

The HTRN was trained in two stages on ImageNet-1k (Deng et al., 2009; Russakovsky et al., 2015). The ResNet-50 backbone was first initialized with pretrained ImageNet weights and frozen, and only the top-down feedback modules were trained; the entire network was then fine-tuned end-to-end using differential learning rates for the backbone and feedback modules.

To isolate the contribution of top-down feedback from other differences between variants, the three representations used in the RSA below were all extracted as different readout stages of the same fine-tuned HTRN: HTRN-FF corresponds to a single feedforward pass with no recurrence, HTRN-LR to the network at full local recurrent depth without top-down feedback, and HTRN-TD to the same full-depth network additionally receiving the accumulated top-down feedback signal.

To align model representations to neural data, feature activations were extracted from the corresponding stages of each of the three HTRN variants (FF, LR, and TD) and mapped onto the three cortical ROIs examined in the MEG analyses. Early network stages (Conv2–3) were taken to correspond to early visual cortex (V1–3), intermediate stages (Conv4) to the LOC, and later stages (Conv5) to IT–PHC. These layer-wise activations were used to construct model RDMs, enabling direct comparison between model representations and time-resolved neural representational structure. This stage-to-region assignment follows the correspondence between successive network stages and successive ventral-stream areas used in previous model–brain comparisons of this pathway (Kietzmann et al., 2019; Rajaei et al., 2019), and was fixed a priori rather than optimized against the neural data.

#### Model behavioral evaluation

Recognition performance was evaluated for every model variant on the same stimulus set used in the MEG experiment (4 object categories × 3 occlusion levels × 64 exemplars = 768 images). For each variant, the output of the final convolutional stage was globally average-pooled to a single feature vector per image (2048 dimensions for the HTRN variants), and a multiclass linear support-vector classifier (LinearSVC; C = 0.1; features standardized on the training split) was trained to discriminate all four object categories jointly, so that chance performance was 25%.

Performance at a given occlusion level was estimated by 8-fold stratified cross-validation over the images of that level, with the training set of every fold additionally containing all images from the remaining occlusion levels; only images of the target occlusion level were ever used for testing. The whole procedure was repeated 20 times with different random fold assignments, and accuracies are reported as the mean across folds and repetitions, with the standard error of the mean across repetitions. Identical feature extraction, classifier and cross-validation settings were used for the HTRN readout stages, for the BLT-VS variants (BLT-FF, BLT-LR and BLT-TD; native configurations B, BL and BLT) and for the CORnet variants (CORnet-Z and CORnet-RT), so that the architectures differ only in the representations entering the classifier.

Human performance is shown in Supplementary Fig. 5 as a reference only. It is taken from the behavioral data of the original MEG-Occlusion experiment (Rajaei et al., 2019), in which observers performed the same four-alternative categorization task; the shaded bands show the reported uncertainty of those estimates.

As a positive control on the feedback implementation, the trained HTRN was evaluated on the full ImageNet-1k validation set (Deng et al., 2009; Russakovsky et al., 2015) alongside the original torchvision ResNet-50 (IMAGENET1K_V2 weights) (He et al., 2015; Paszke et al., 2019) used to initialize its backbone, both scored with the identical pipeline used throughout. Because the two fixed classifiers were scored on the same images, paired per-image top-1 and top-5 correctness were compared with McNemar’s test (McNemar, 1947), with a nonparametric bootstrap (100,000 resamples) on the accuracy difference as the accompanying effect-size estimate.

#### Significance analysis

Statistical significance for time-resolved analyses was evaluated using nonparametric cluster-based permutation tests (Maris & Oostenveld, 2007) as implemented in MNE-Python (Gramfort et al., 2013). For each test, a one-sample t test against the null value was computed at every time point across the 14 participants; the null value was 0.5 for decoding, zero for the RSA correlations, the prestimulus baseline for Granger causality, and zero for every difference between conditions, ROIs or model variants. Time points whose t value exceeded the parametric threshold corresponding to *p* < 0.05 with 13 degrees of freedom were grouped into contiguous clusters, and each cluster was summarized by the sum of its t values.

Cluster-level significance was assessed against the exact permutation distribution obtained by enumerating all sign flips of the participant-level data, and clusters with *p* < 0.05 were considered significant. Tests of above-chance decoding, of differences between conditions, ROIs or model variants, and of within–cross decoding differences were two-tailed; tests of Granger causality against the prestimulus baseline were one-tailed, because only increases in predictability are interpretable. Clustering was performed over −100 to 600 ms for the decoding and representational similarity analyses and over 0 to 300 ms for the Granger-causality analyses.

Onset and peak latencies were estimated with a jackknife procedure: the group-average time course was recomputed for each of the 14 leave-one-participant-out subsamples, and the peak of that subsample (for onsets, the first time point of its earliest significant cluster) was extracted. Reported latencies are the mean and standard deviation across the 14 subsamples.

Cluster-based correction controls the family-wise error rate within a single time course but not across the several families of tests reported here (3 ROIs × 2 occlusion levels for decoding, 6 directed ROI pairs × 2 occlusion levels for Granger causality, and 3 ROIs × 2 occlusion levels × 3 variant contrasts for the representational similarity analysis). Cluster p-values are therefore reported without correction across families, and individual clusters should be read as descriptive; the conclusions drawn here rest on the convergence of the decoding, masking, connectivity, and modeling analyses rather than on any single cluster.

## Supporting information

**Supplementary Figure 1.**
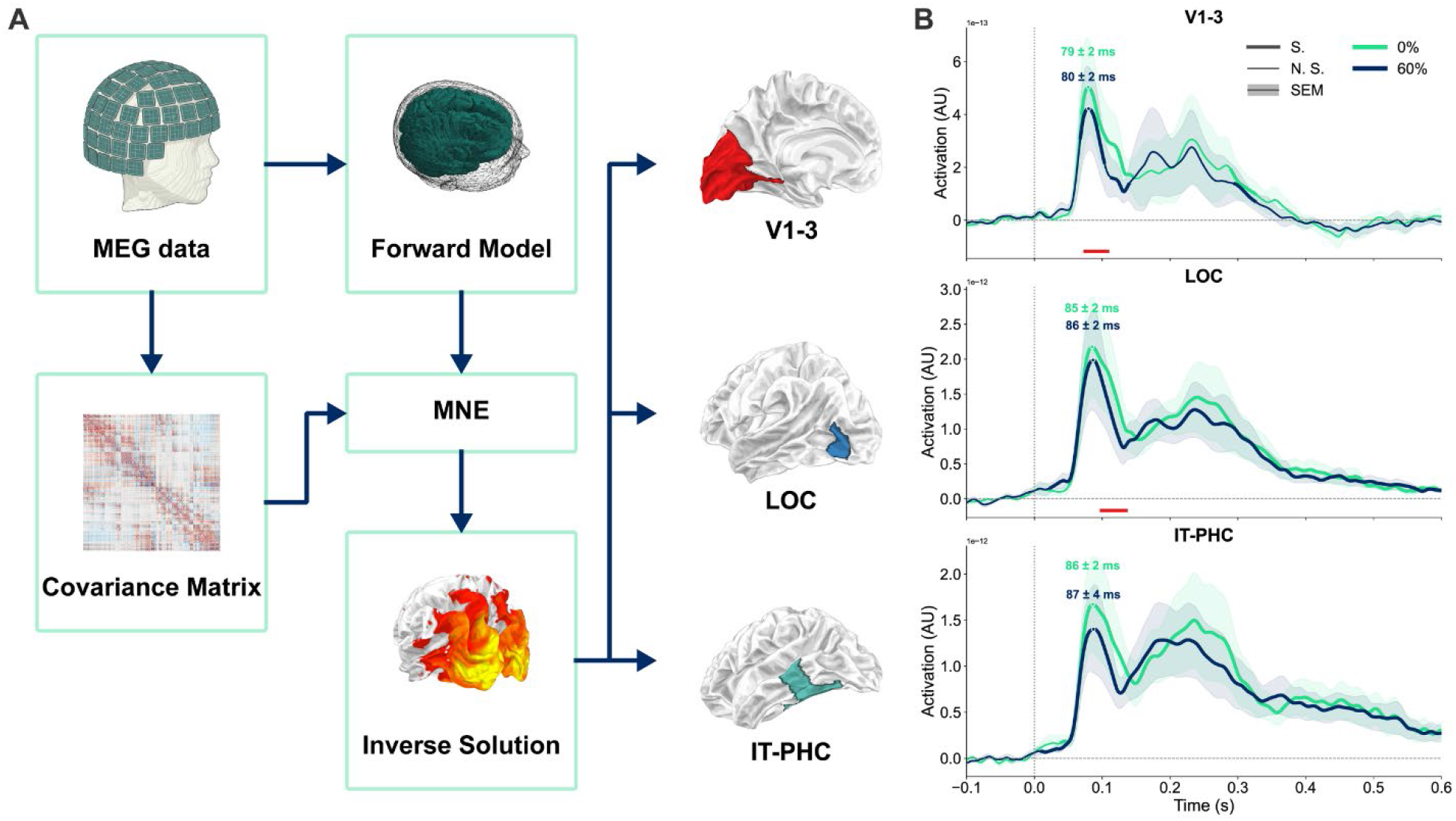
Source-estimated evoked responses across ventral visual ROIs. **A)** Schematic overview of source reconstruction and ROI time-course extraction. Preprocessed MEG data, the forward model, and the noise covariance matrix were used to construct minimum-norm inverse operators. Source-space activity was then summarized within three predefined ROIs: V1–3, LOC, and IT– PHC. **B)** Source-estimated evoked responses in V1–3, LOC, and IT–PHC under 0% and 60% occlusion (N = 14). Curves show the mean across participants, and shaded areas indicate the standard error of the mean (SEM). Labels indicate peak response latencies (mean ± SD). Red horizontal bars indicate time intervals in which evoked-response amplitude differed significantly between occlusion conditions (cluster-based permutation tests, *p* < 0.05). Stimulus onset occurred at 0 s.

**Supplementary Figure 2.**
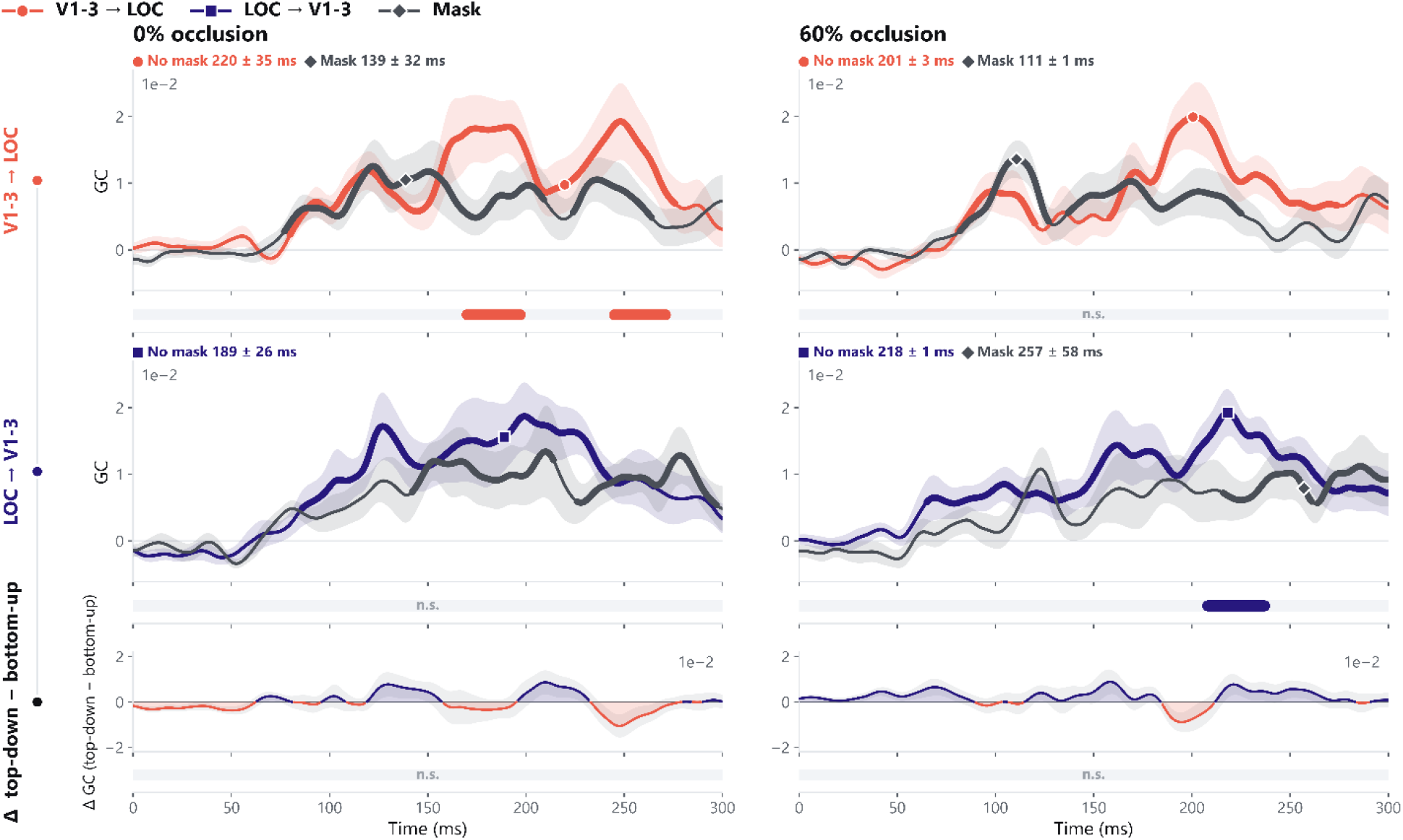
Directional representational Granger causality between V1–3 and LOC. Grand-average representational Granger causality time courses for the two directions of information flow between V1–3 and LOC, shown separately for the no-mask (colored lines) and mask (dark gray lines) conditions (N = 14). Rows correspond to bottom-up flow (V1–3 → LOC), top-down flow (LOC → V1–3), and the difference between the two directions (Δ GC, top-down minus bottom-up); columns correspond to the 0% and 60% occlusion conditions. Granger causality values are expressed relative to the prestimulus baseline. Shaded areas indicate the standard error of the mean (SEM). Thickened portions of the curves mark time periods with significant above-baseline information flow, and labels above the curves denote peak latencies (mean ± SD). Horizontal bars beneath each panel indicate time windows of a significant difference between the no-mask and mask conditions (cluster-based permutation test, *p* < 0.05), colored by the dominant condition; “n.s.” denotes the absence of a significant cluster. In the bottom row, the area between the curve and zero is filled in the top-down color above zero and in the bottom-up color below zero, so that the dominant direction at each time point is read directly from the fill.

**Supplementary Figure 3.**
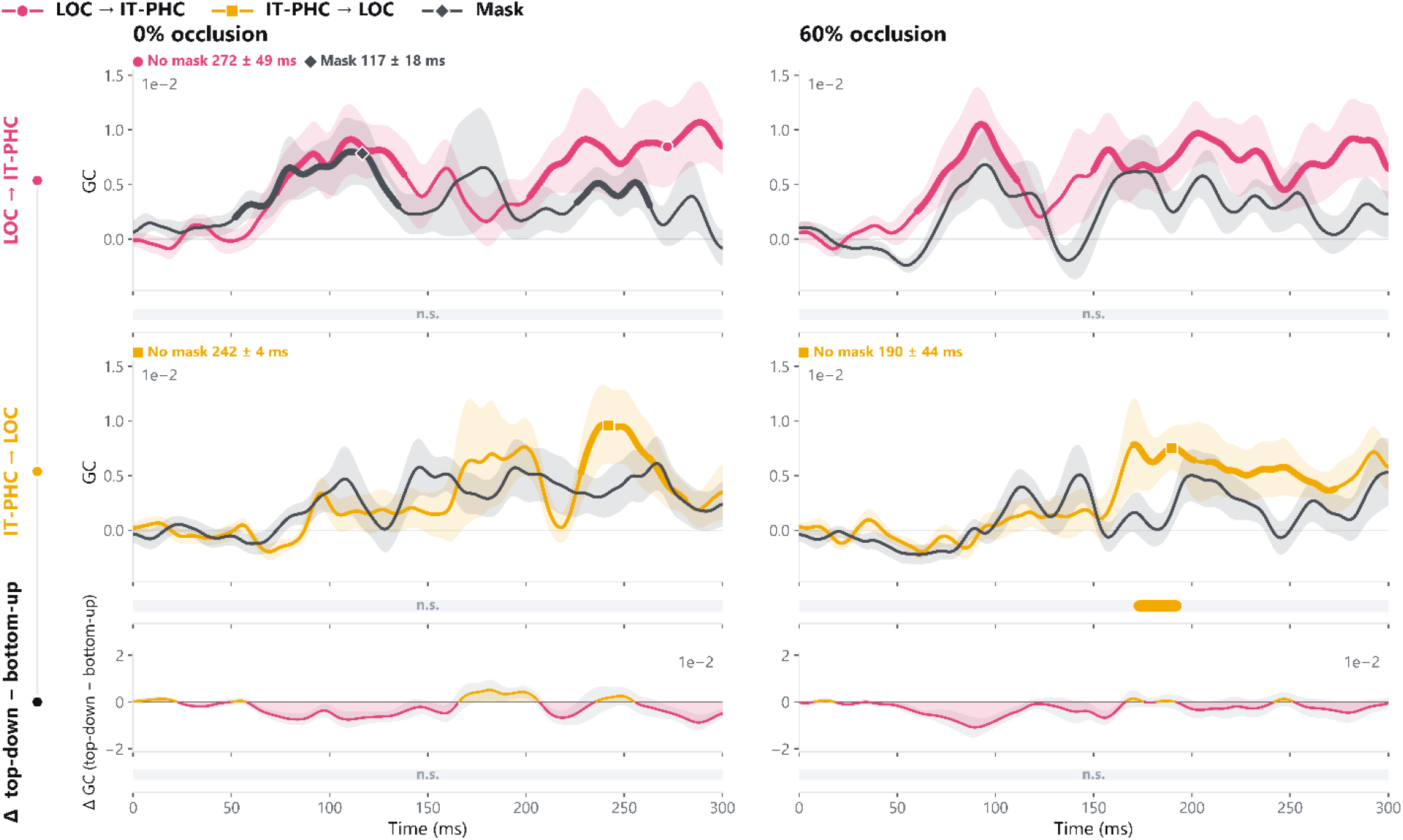
Directional representational Granger causality between LOC and IT–PHC. Grand-average representational Granger causality time courses for the two directions of information flow between LOC and IT–PHC, shown separately for the no-mask (colored lines) and mask (dark gray lines) conditions (N = 14). Rows correspond to bottom-up flow (LOC → IT–PHC), top-down flow (IT–PHC → LOC), and the difference between the two directions (Δ GC, top-down minus bottom-up); columns correspond to the 0% and 60% occlusion conditions. Granger causality values are expressed relative to the prestimulus baseline. Shaded areas indicate the standard error of the mean (SEM). Thickened portions of the curves mark time periods with significant above-baseline information flow, and labels above the curves denote peak latencies (mean ± SD). Horizontal bars beneath each panel indicate time windows of a significant difference between the no-mask and mask conditions (cluster-based permutation test, *p* < 0.05), colored by the dominant condition; “n.s.” denotes the absence of a significant cluster. Conventions are otherwise as in Supplementary Figure 2.

**Supplementary Figure 4.**
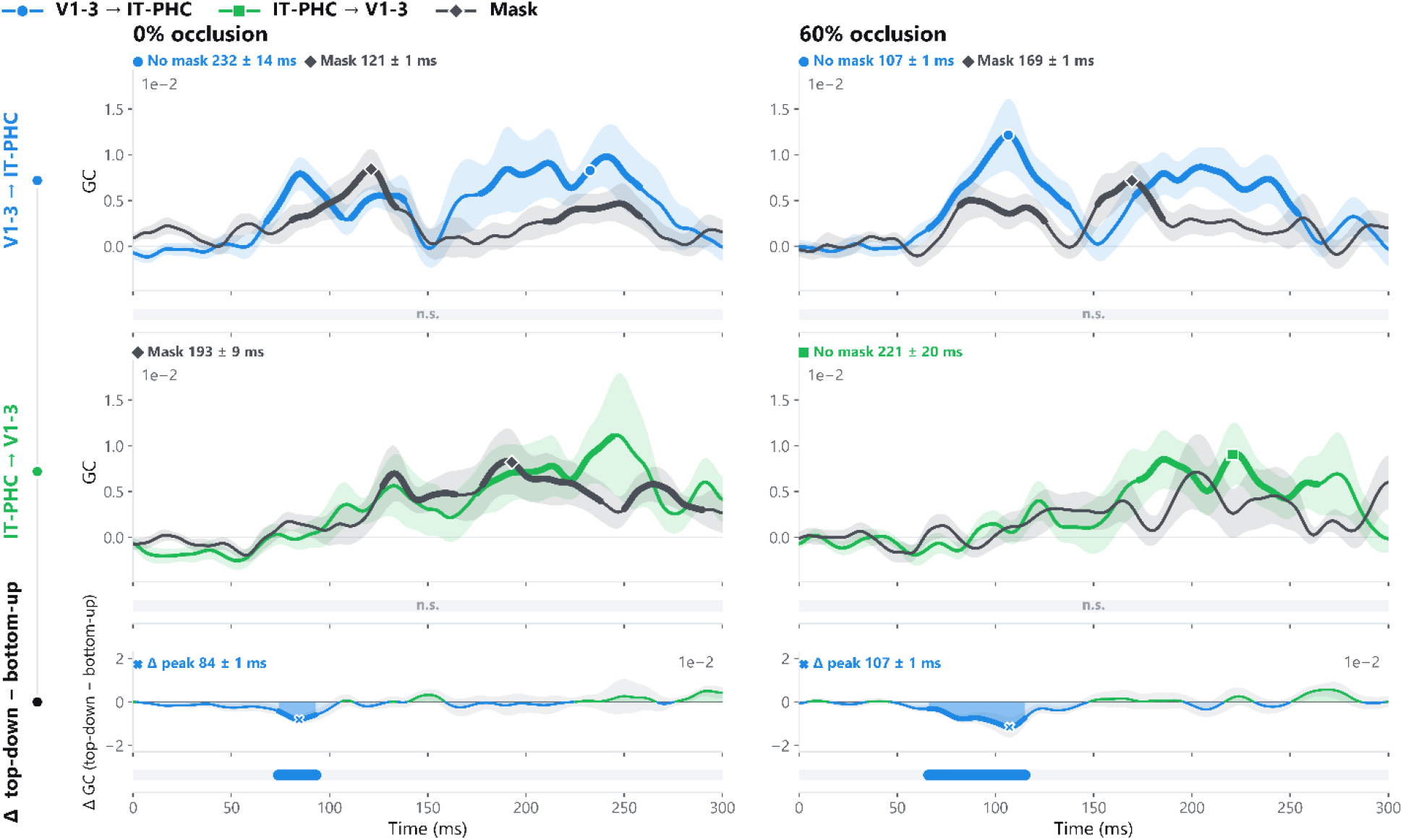
Directional representational Granger causality between V1–3 and IT–PHC. Grand-average representational Granger causality time courses for the two directions of information flow between V1–3 and IT–PHC, shown separately for the no-mask (colored lines) and mask (dark gray lines) conditions (N = 14). Rows correspond to bottom-up flow (V1–3 → IT–PHC), top-down flow (IT–PHC → V1–3), and the difference between the two directions (Δ GC, top-down minus bottom-up); columns correspond to the 0% and 60% occlusion conditions. Granger causality values are expressed relative to the prestimulus baseline. Shaded areas indicate the standard error of the mean (SEM). Thickened portions of the curves mark time periods with significant above-baseline information flow, and labels above the curves denote peak latencies (mean ± SD). Horizontal bars beneath each panel indicate time windows of a significant difference between the no-mask and mask conditions (cluster-based permutation test, *p* < 0.05), colored by the dominant condition; “n.s.” denotes the absence of a significant cluster. Conventions are otherwise as in Supplementary Figure 2.

**Supplementary Figure 5.**
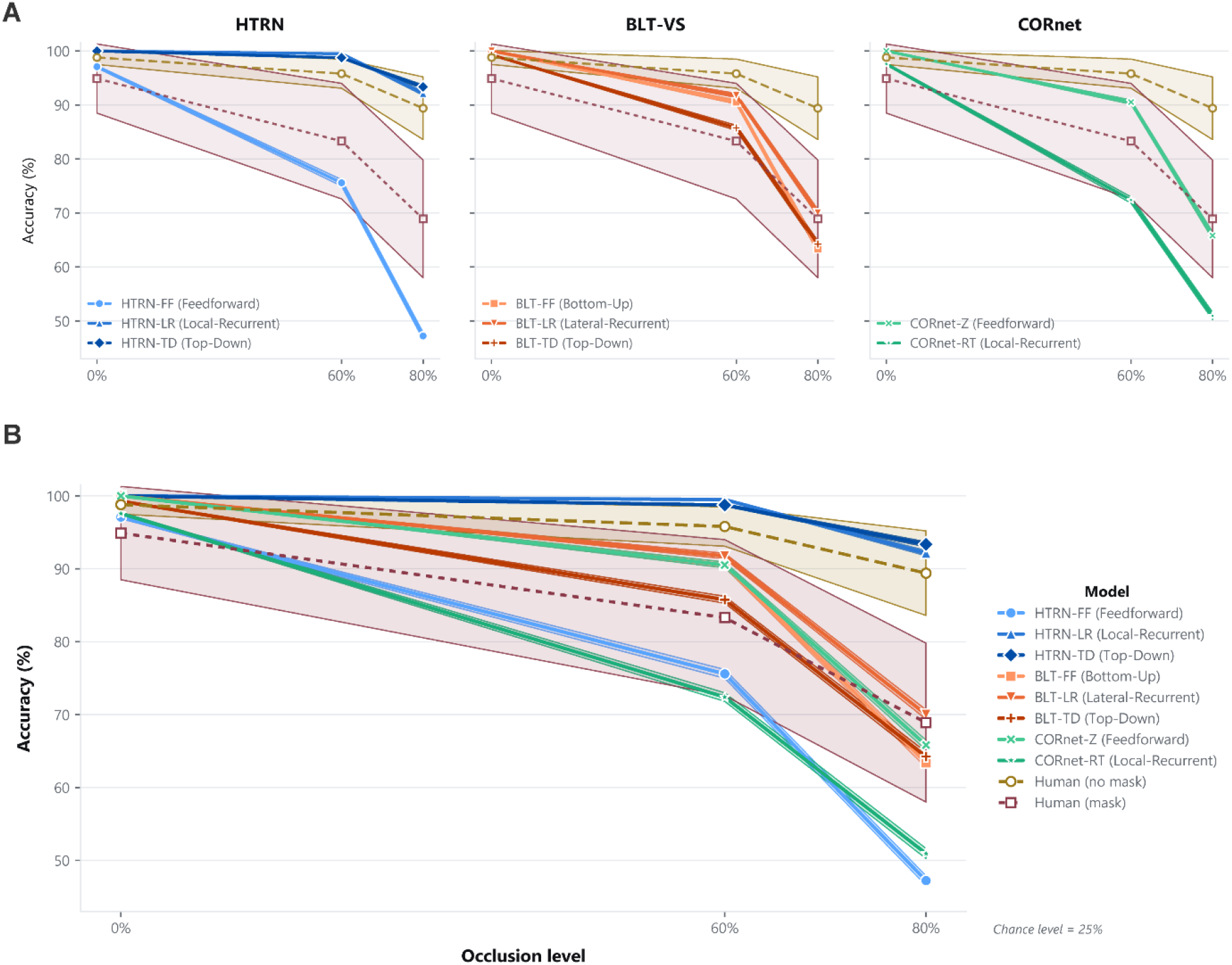
Recognition robustness across recurrent model architectures. Four-category classification accuracy for HTRN, BLT-VS, and CORnet variants under 0%, 60%, and 80% occlusion. The upper panels show performance separately within each architecture; the lower panel shows all model variants together. Solid colored lines represent model accuracy. Dashed lines and shaded bands show human performance and its uncertainty in the no-mask and mask conditions. Chance performance was 25%. All models performed near ceiling under unoccluded viewing, but their performance diverged as occlusion increased. HTRN-LR and HTRN-TD remained closest to human performance at high occlusion, whereas HTRN-FF and the BLT-VS and CORnet variants showed substantially larger declines (behavioral accuracies adopted from Rajaei et al., 2019).

**Supplementary Figure 6.**
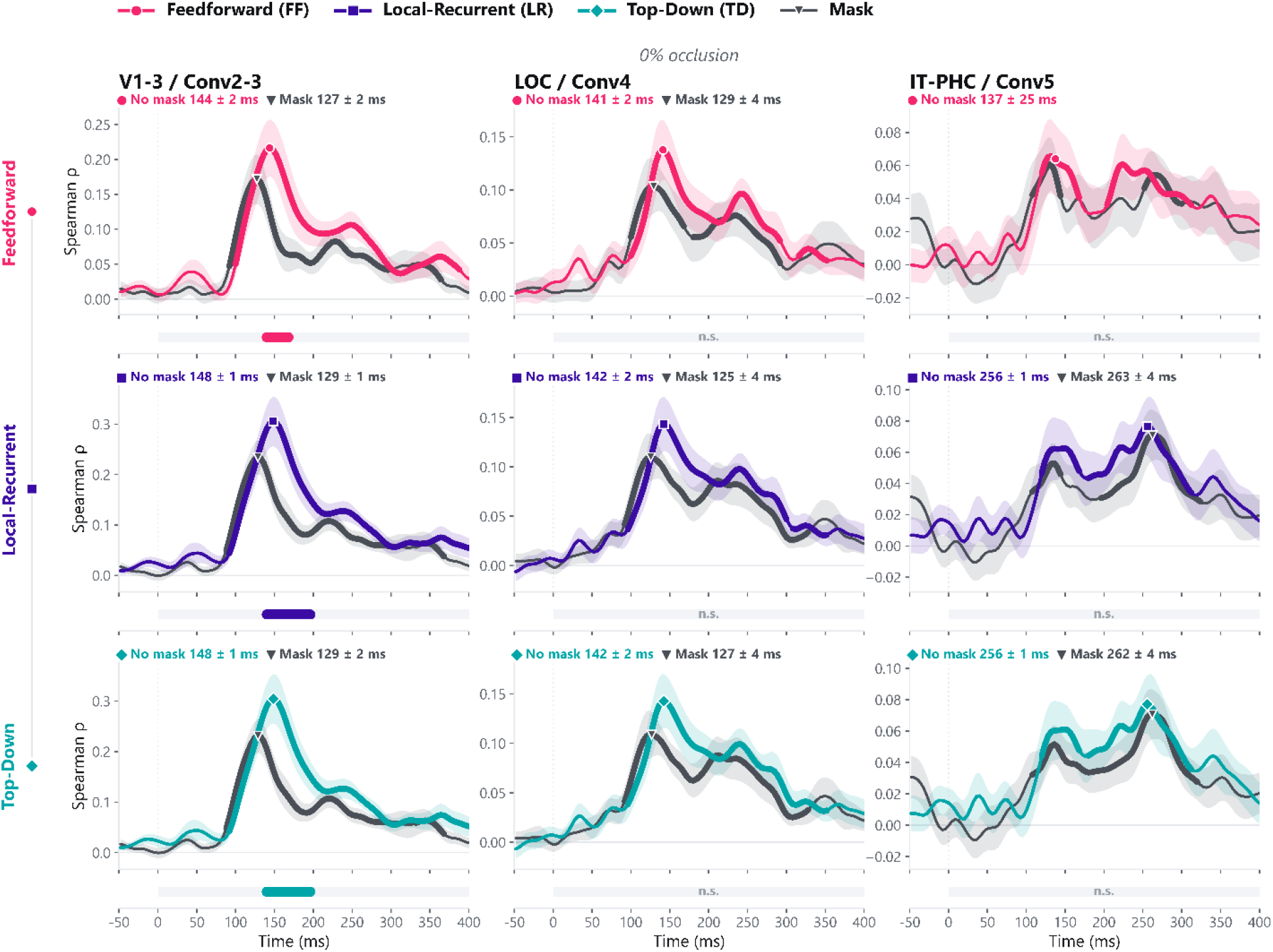
Masking effects on model–brain correspondence without occlusion. Time-resolved Spearman correlations between human neural and HTRN representational geometries for no-mask and mask trials under 0% occlusion (N = 14). Rows show the FF, LR, and TD variants; columns show the V1–3/Conv2–3, LOC/Conv4, and IT–PHC/Conv5 mappings. Colored curves indicate no-mask trials and gray curves indicate masked trials. Shaded areas show the SEM. Thickened segments indicate significant correlations greater than zero; labels show peak correlation latencies (mean ± SD). Horizontal bars mark significant differences between conditions (cluster-based permutation tests, *p* < 0.05), with colors indicating the condition with greater correspondence. “n.s.” indicates no significant cluster.

**Supplementary Figure 7.**
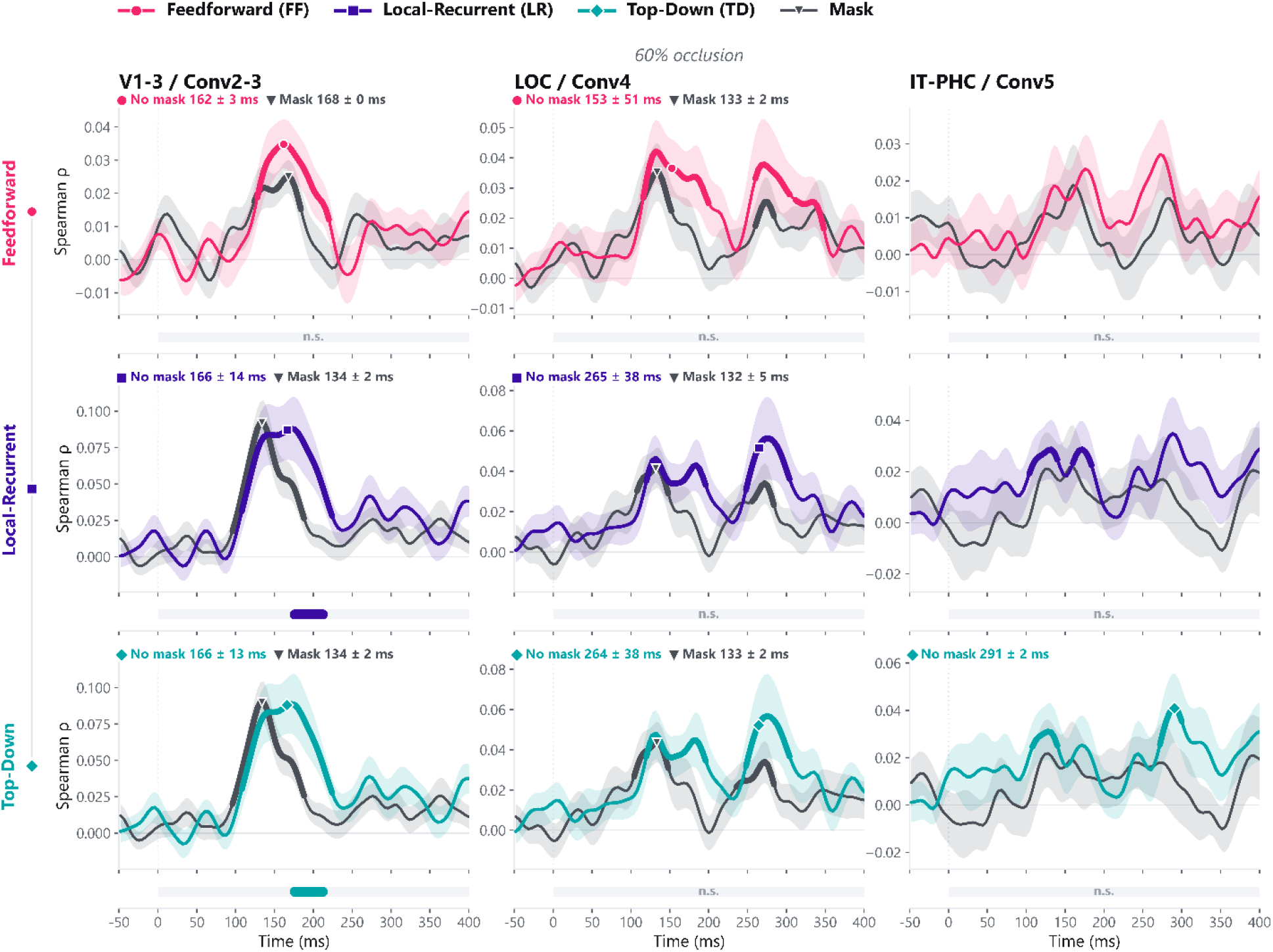
Masking effects on model–brain correspondence under occlusion. Same as Supplementary Fig. 6, but for 60% occlusion. Model–brain correspondence is shown for no-mask and mask trials for the FF, LR, and TD variants of the HTRN across the V1–3/Conv2–3, LOC/Conv4, and IT–PHC/Conv5 mappings.

**Supplementary Figure 8.**
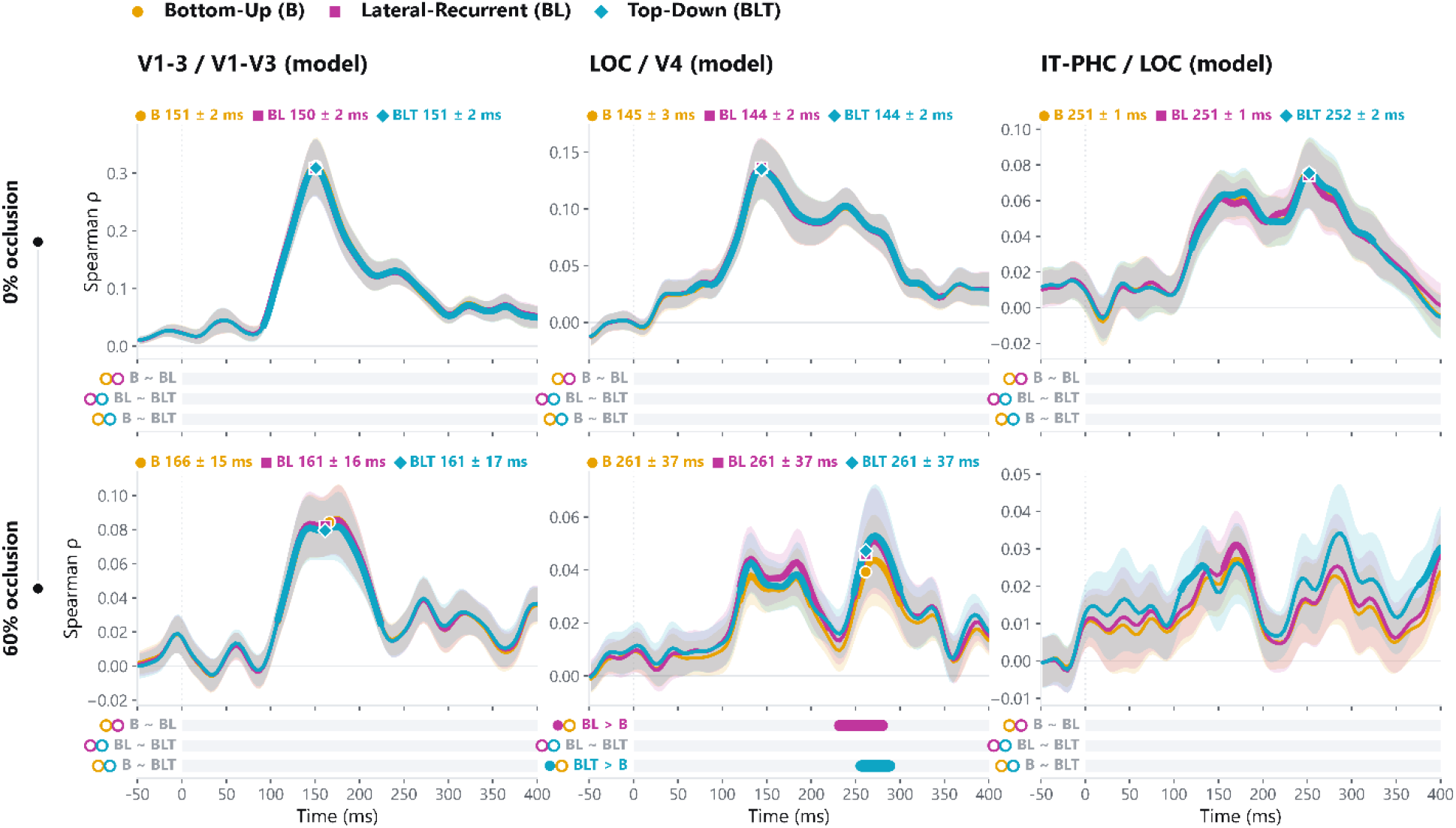
BLT model–brain correspondence without masking. Time-resolved Spearman correlations between human neural representational geometry and three BLT-VS variants under 0% and 60% occlusion without masking (N = 14): a feedforward variant (BLT-FF; native configuration B), a variant with local recurrent connections (BLT-LR; BL), and a variant with local recurrence plus top-down feedback (BLT-TD; BLT). Columns show the V1–3/V1–V3, LOC/V4, and IT– PHC/LOC mappings. Curves show the mean across participants; shaded areas indicate the SEM. Thickened segments indicate significant correlations greater than zero, and labels show peak latencies (mean ± SD). Horizontal bars mark significant pairwise differences between variants (cluster-based permutation tests, *p* < 0.05), with colors indicating the variant with greater correspondence. “n.s.” indicates no significant cluster.

**Supplementary Figure 9.**
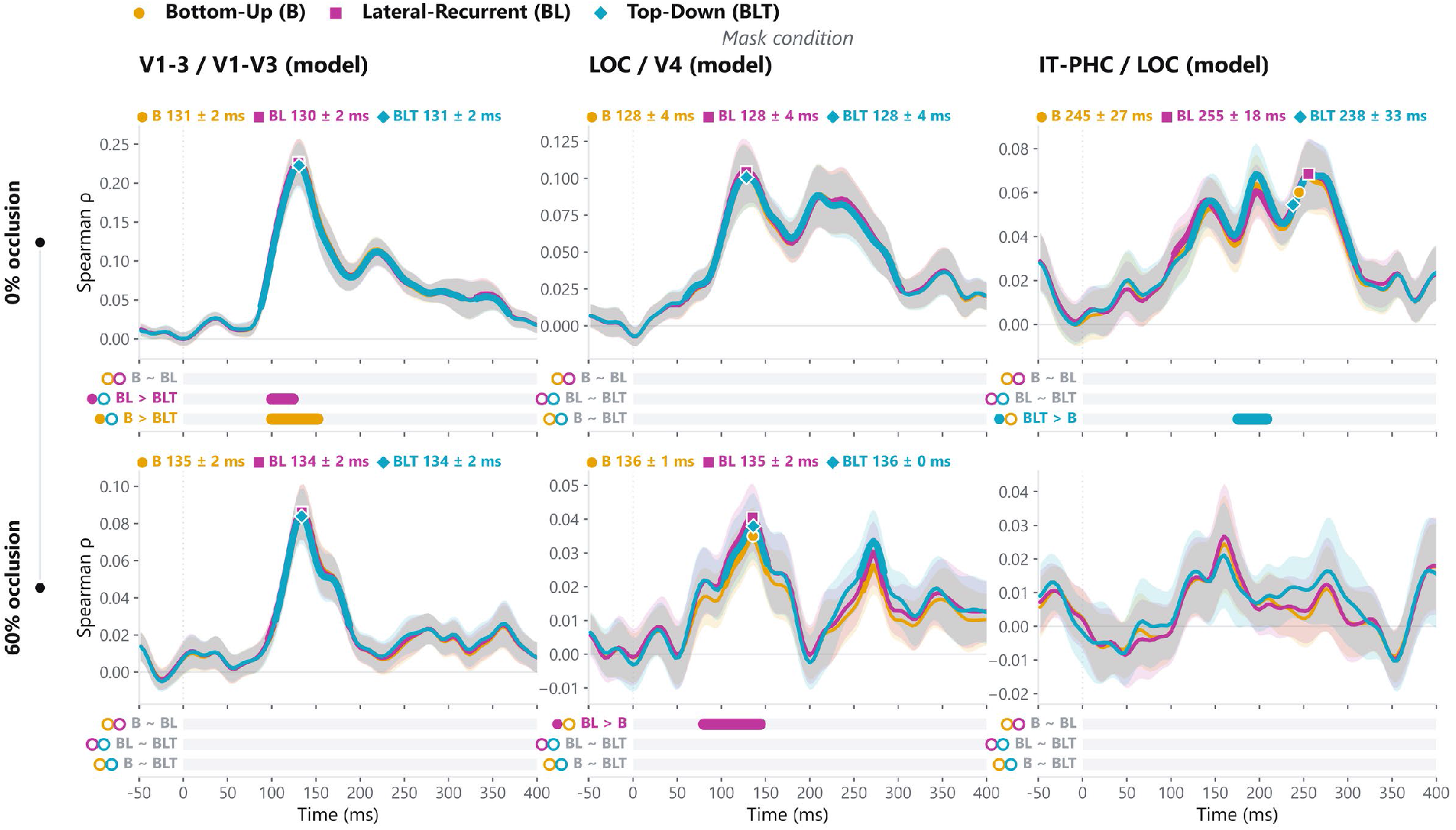
BLT model–brain correspondence on masked trials. Same analysis as Supplementary Fig. 8, but computed on trials with a backward mask. Results are shown for 0% and 60% occlusion across the V1–3/V1–V3, LOC/V4, and IT–PHC/LOC mappings. Conventions otherwise follow Supplementary Fig. 8.

**Supplementary Figure 10.**
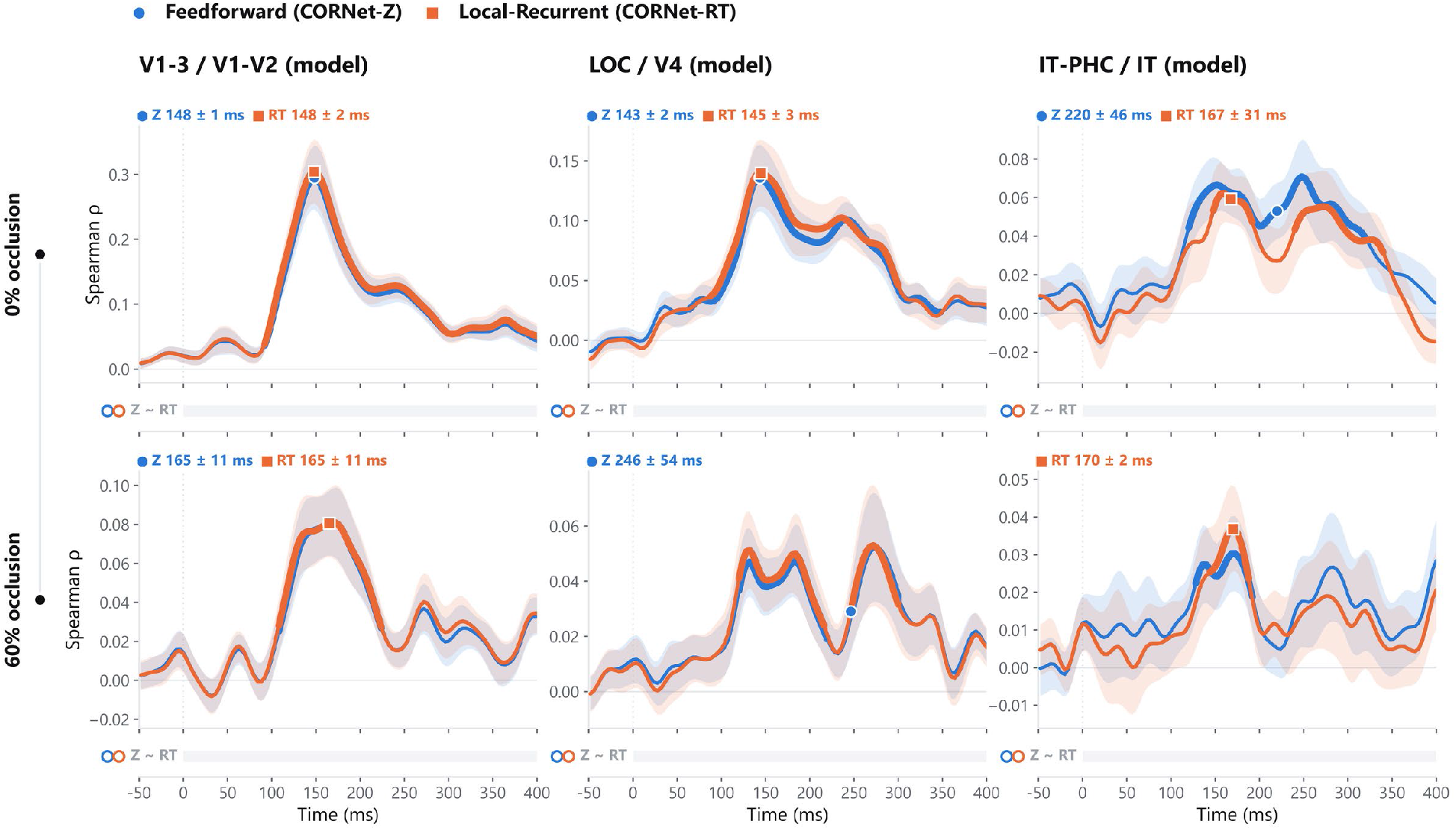
CORnet model–brain correspondence without masking. Time-resolved Spearman correlations between human neural representational geometry and CORnet-Z, a feedforward model, and CORnet-RT, a model with local recurrence (Kubilius et al., 2019), under 0% and 60% occlusion without masking (N = 14). Columns show the V1–3/V1–V2, LOC/V4, and IT–PHC/IT mappings. Curves show the mean across participants; shaded areas indicate the SEM. Thickened segments indicate significant correlations greater than zero, and labels show peak latencies (mean ± SD). Horizontal bars indicate significant pairwise differences between models (cluster-based permutation tests, *p* < 0.05), with colors denoting the model with greater correspondence. “n.s.” indicates no significant cluster.

**Supplementary Figure 11.**
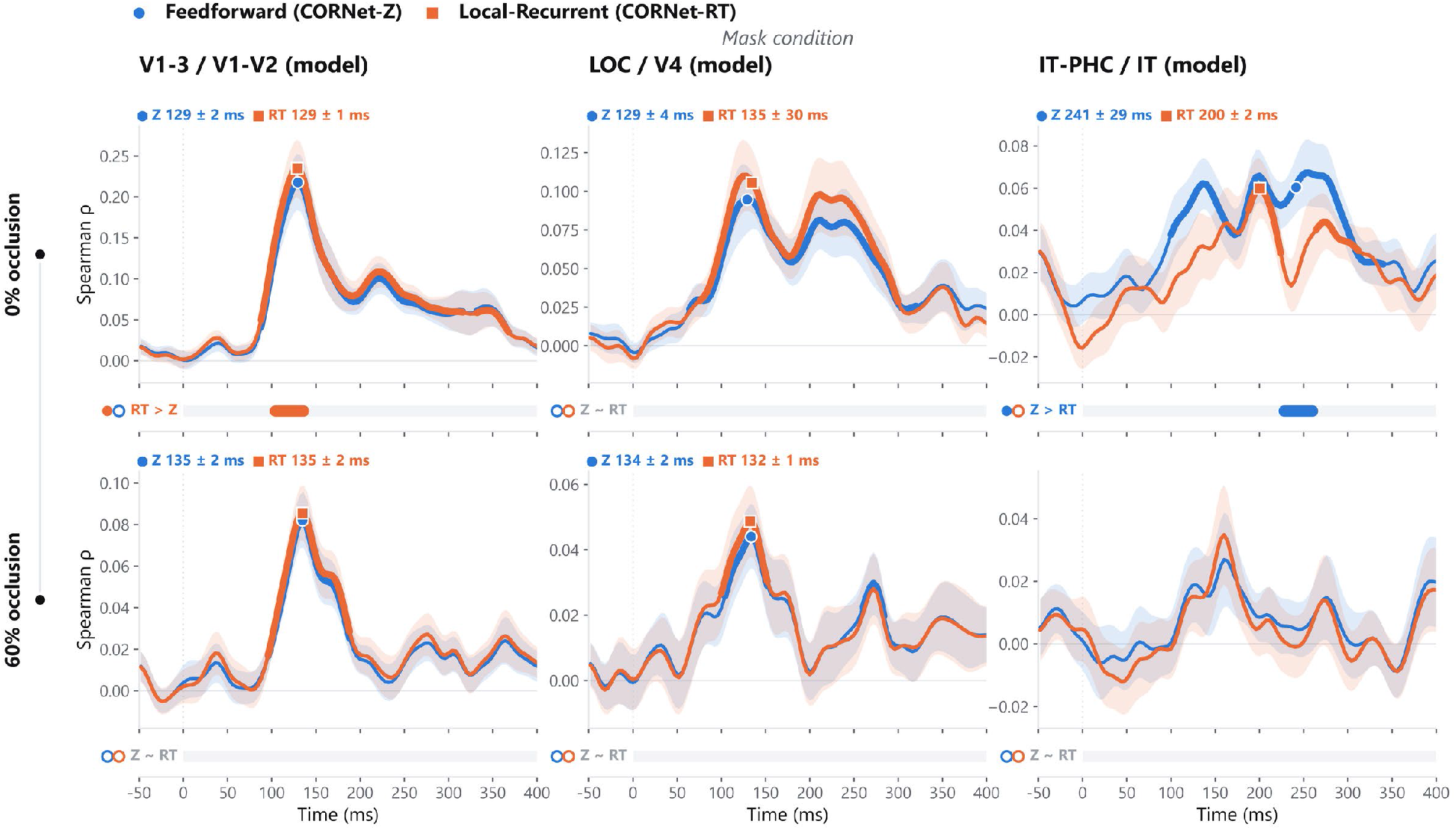
CORnet model–brain correspondence on masked trials. Same analysis as Supplementary Fig. 10, but computed on trials with a backward mask. Results are shown for 0% and 60% occlusion across the V1–3/V1–V2, LOC/V4, and IT–PHC/IT mappings. Conventions otherwise follow Supplementary Fig. 10

